# Placental *hIGF1* nanoparticle gene therapy in guinea pigs ameliorates fetal growth restriction-associated changes in hepatic lipid and glucose metabolism-related signaling pathways in a fetal sex-specific manner

**DOI:** 10.1101/2024.10.30.621100

**Authors:** Baylea N Davenport, Alyssa Williams, Timothy RH Regnault, Helen N Jones, Rebecca L Wilson

## Abstract

Fetal development in an adverse in utero environment significantly increases the risk of developing metabolic diseases in later life, including dyslipidemia, non-alcoholic fatty liver diseases and diabetes. The aim of this study was to determine whether improving the in utero fetal growth environment with a placental nanoparticle gene therapy would ameliorate fetal growth restriction (FGR)-associated dysregulation of fetal hepatic lipid and glucose metabolism-related signaling pathways. Using the guinea pig maternal nutrient restriction (MNR) model of placental insufficiency and FGR, placenta efficiency was significantly improved following three administrations of a non-viral polymer-based nanoparticle gene therapy to the placenta from mid-pregnancy (gestational day 35) until gestational day 52. The nanoparticle gene therapy transiently increased expression of *human insulin-like growth factor 1* (*hIGF1*) in placenta trophoblast. Fetal liver tissue was collected near-term at gestational day 60. Fetal sex specific differences in liver gene and protein expression of pro-fibrosis and glucose metabolism-related factors were demonstrated in sham-treated FGR fetuses but not observed in FGR fetuses who received placental *hIGF1* nanoparticle treatment. Increased plasma bilirubin, and indirect measure of hepatic activity, was also increased with placental *hIGF1* nanoparticle treatment. We speculate that the changes in liver gene and protein expression and increased liver activity that result in similar expression profiles to appropriately growing Control fetuses might confer protection against increased susceptibility to aberrant liver physiology in later-life. Overall, this work opens avenues for future research assessing the translational prospect of mitigating FGR-induced metabolic derangements.

**NEW AND NOTEWORTHY:** A placenta-specific non-viral polymer-based nanoparticle gene therapy that improves placenta nutrient transport and near-term fetal weight ameliorates growth restriction-associated changes to fetal liver activity, and cholesterol and glucose/nutrient homeostasis genes/proteins that might confer protection against increased susceptibility to aberrant liver physiology in later-life. This knowledge may have implications towards removing predispositions that increase the risk of metabolic diseases including diabetes, dyslipidemia and non-alcoholic fatty liver disease in later life.

## INTRODUCTION

Fetal growth restriction (FGR) is a condition characterized by failure of the fetus to meet their genetic growth potential during pregnancy, often due to placental insufficiency and/or maternal factors (1). This suboptimal intrauterine environment can initiate a cascade of developmental adaptations, collectively referred to as developmental programming, which have immediate and long-lasting effects on an individual’s health. One significant aspect of this programming is epigenetics, a process whereby gene expression is regulated through modifications which do not change the DNA sequence but can stably alter gene activity and is influenced by environmental factors during critical periods of development. The liver is a critical organ for metabolic regulation that early in utero development and growth is critical for successful later life metabolic functioning (2). Altered liver development and function is observed in FGR, and it is understood epigenetic modifications underly these changes (3, 4), leading to altered expression of genes and proteins involved in lipid and glucose homeostasis (5). This reprogramming, while understood to initially confer a survival strategy in utero, appears to predispose individuals to metabolic diseases after birth (postnatally) such as dyslipidemia, non-alcoholic fatty liver disease (NAFLD), insulin resistance, obesity and diabetes later in life (6). Thus, the developmental origins of these metabolic disorders are closely linked to the adaptive responses triggered by FGR, highlighting the crucial interplay between early development and growth conditions and long-term liver function as it relates to metabolic health.

Physiologically the liver functions to convert excess carbohydrates to lipids (fatty acids and triglycerides) (7) and regulate cholesterol homeostasis through uptake, export, biosynthesis and storage (8). However, disruptions to hepatic lipid metabolism-related pathways, for example through excess lipid consumption and/or genetic susceptibility, result in lipid accumulation and a lipotoxic state in the liver (9, 10) ultimately contributing to increased susceptibility to metabolic diseases later in life. Prior studies have shown that placental insufficiency and the resulting low birth weight of the offspring is associated with microvesicular hepatic steatosis (11), programmed hepatic cholesterol accumulation and impaired cholesterol elimination (12). Compromised lipid metabolism and elimination occurs in parallel to increased liver expression of pro-fibrotic and inflammatory genes (11) suggesting a combination of molecular signaling events that ultimately lead to dysregulated hepatic lipid homeostasis that is associated with FGR.

The liver is also a major organ important for insulin action and glucose homeostasis including gluconeogenesis and nutrient-sensing (13). The actions of insulin in the liver are mediated by PI3K/AKT signaling pathways that in turn regulated the expression of glucose metabolism-related genes such as gluconeogenesis enzymes, growth factors and glucose transporters to ultimately increase glucose production (14). Regulation of these glucose metabolism-related pathways is highly sensitive to changes in nutrient status and cell stress (15). Prior studies in the near term fetus have demonstrated increased hepatic expression of glucose metabolism-related factors (16, 17) which persist postnatally, likely contributing to increased adiposity, insulin resistance and glucose intolerance (18, 19). These outcomes were more prevalent in male offspring compared to female highlighting a nuanced understanding of how early-life conditions can predispose individuals to metabolic syndromes based on fetal sex. Collectively, these studies contribute to a growing body of evidence on how FGR conditions and early-life nutrition can significantly influence liver health and the risk of developing metabolic disorders which may be mitigated with effective therapeutics that improve the in utero environment and fetal growth.

Observational human studies have provided understanding of the relationship between FGR and developmental programming of adverse long-term health and are supported by mechanistic studies that rely on pre-clinical models including animal model systems. Pregnant guinea pigs are often used to study the mechanisms linking FGR and developmental programming of the liver because of their unique reproductive physiological and fetoplacental developmental characteristics that are similar to human reproductive physiology. Characteristics include similar placenta structure and function (20), extended gestational developmental that results in offspring being born precocial (21), and the metabolic and nutritional physiology of guinea pigs is more like humans compared to other rodents (22, 23). Additionally, guinea pigs exhibit a natural predisposition to develop non-communicable diseases, including obesity, atherosclerosis, and diabetes, making them particularly valuable for studies investigating the long-term effects of FGR on the development of metabolic syndromes and liver diseases (24).

In the guinea pig maternal nutrient restriction (MNR) model of FGR we have demonstrated repeated treatment of the placenta with a polymer-based nanoparticle for non-viral, transient human *Insulin-like 1 Growth Factor* (*hIGF1*) gene delivery (25) across the second half of pregnancy to near term results in improved placenta efficiency, increased fetal blood glucose, reduced fetal blood cortisol and improved fetal weight when compared to sham treated FGR fetuses (26). IGF1 is a master regulator of placental development and function, regulating cell proliferation/ differentiation, vasculature structure, nutrient transport, and hormone production, and is downregulated in human FGR (27, 28). Previous animal-based studies have assessed providing IGF1 to the fetus, amniotic fluid or placenta for the treatment of FGR with varied outcomes (29–31) and not comprehensively assessed the implications to fetal organ development and physiology. We hypothesize that improving placenta function can directly increase the nutrient availability to the fetus without necessitating the fetal liver to metabolize and synthesize nutrients from a limited supply. This approach could potentially alleviate the metabolic burden on the developing fetal liver in situations of FGR, ensuring more efficient growth and development by optimizing the nutrient and oxygen availability directly from the placental circulation (32, 33). Our aim was to assess fetal liver growth and activity, lipid composition and expression of genes and proteins involved in lipid and glucose metabolism-related signaling pathways following repeated placenta *hIGF1* nanoparticle gene therapy which improved fetal growth trajectories.

## MATERIALS & METHODS

### Nanoparticle Formation

Detailed methods of copolymer synthesis and nanoparticle formation have been previously published (25). Briefly, lyophilized PHPMA_115_-b-PDMEAMA_115_ co-polymer was reconstituted in sterile saline, complexed with 50 µg of plasmid containing the *hIGF1* gene and *CYP19A1* promotor under aseptic conditions, and stored at -20°C until administration.

### Animal Husbandry and Nanoparticle Delivery

Animal care and usage was approved by the Institutional Animal Care and Usage Committee at the University of Florida (Protocol #202011236). Detailed methods of Animal Husbandry and Nanoparticle Delivery is previously published (26). Dams (Dunkin-Hartley) were purchased at 500-550 g (8-9 weeks of age) and allowed to acclimate to the animal facilities for two weeks prior to study initiation. After the acclimatization period, dams were weighed, ranked from heaviest to lightest and systematically assigned to either ad libitum diet (termed Control: n = 6) or maternal nutrient restriction (MNR) diet (n=12) so that the mean weight was no different between the Control and MNR groups. For both Control and MNR dams, water was provided ad libitum. Food (LabDiet diet 5025: 27% protein, 13.5% fat, and 60% carbohydrate as % of energy) intake in the MNR dams was however, restricted to 70% per kilogram body weight of the Control group from at least 4 weeks preconception through to mid-pregnancy (GD30). From GD30, food was restricted to 90% per kilogram body weight of the Control group to prevent fetal demise and pregnancy loss (34). Control dams were provided food ad libitum.

At gestational day (GD) 36±3, dams were anesthetized, and an ultrasound scan determined the number and location of fetuses. Typically, guinea pigs carry three fetuses, asymmetrically positioned so that one uterine horn contains one fetus, whilst the other uterine horn contains two fetuses. The uterine horn containing one fetus was selected, and nanoparticle was delivered to the placenta via ultrasound-guided intra-placental injection. Only one placenta per litter was injected with either a non-expressing sham nanoparticle (n=12; 6 Control dams and 6 MNR dams) or *hIGF1* nanoparticle (n=6 MNR + *hIGF1* dams). Fetuses whose placentas received the injection were designated the “Directly Treated” fetus. All remaining fetuses in the litter were designated “Indirectly Exposed”. Intra-placental *hIGF1* or sham nanoparticle injections into the same placenta were repeated twice, 8 days apart (GD 44±3 and GD52±3) with all dams successfully receiving three injections. Eight days following the third injection (GD60±3), dams and fetuses were sacrificed by carbon dioxide asphyxiation followed by exsanguination and removal of the heart. Fetal liver tissue (Control: n=8 female and n=11 male, MNR: n=5 female and 11 male, MNR + *hIGF1*: n=6 female and 10 male) was weighed and processed for histology (fixed in 4% PFA), RNA extractions (frozen in RNAlater) and flash-frozen in liquid nitrogen. To ensure rigor and reproducibility, all samples were analyzed in a blinded manner.

### Histology and Immunohistochemistry

Paraffin embedded fetal liver tissue sections (3-5 µm thick) were obtained and liver morphology assessed using histology and immunohistochemistry. Sections were dewaxed and rehydrated following standard protocols. Portal triad (hepatic artery, portal vein and bile duct) morphology and liver lipid accumulation was assessed using hematoxylin and eosin staining. Formal quantification of lipid accumulation was not performed as very few lipid droplets were observed. Portal triad collagen deposition was visualized using Sirius Red staining, imaged on the Axioscan Scanning Microscope (*Zeiss*) and analyzed using ImageJ software (v1.53a). Images (10X magnification) of 10 separate portal triads per section were obtained using the Zen Imaging software (*Zeiss*). The Color Deconvolution plugin in ImageJ was used to split images into red, green and blue channels. With the blue channel image selected, percent area was measured using the Threshold and Measure tools. An average percent collagen was then calculated across the 10 images per section.

For analysis of hepatocyte proliferation and immune cell infiltration, immunohistochemistry was used to visualize Ki67 positive nuclei and CD45 positive immune cells, respectively. Antigen retrieval was performed by incubating slides in 10X Target Retrieval Solution (*Invitrogen*) for 20 min in a bead bath at 95°C and then cooled at room temperature. Endogenous peroxidase activity was suppressed by incubating the slides in 3% hydrogen peroxide and then animal-free protein block (*Vector)* was applied to the sections for 30 min at room temperature. Primary antibodies for Ki67 (*Invitrogen* MA5-14520; diluted 1:200) and CD45 (*Biorad* MCA1130; dilute 1:100) were diluted in the protein block and applied overnight at 4 °C under humidified conditions. Negative controls were included by omitting the primary antibody from the diluent. Ki67 and CD45 antibody binding was amplified with goat anti-rabbit secondary antibody (*Invitrogen*; diluted 1:200) or goat anti-mouse antibody (*Invitrogen*; diluted 1:200), respectively, followed by incubation with ABC reagents (*Vector*) and visualized using 3,3′-diaminobenzidine (DAB) (*Vector*). Nuclei were counterstained with hematoxylin and sections mounted under coverslips using DPX mounting medium (*Millipore*). Staining was imaged on the Axioscan Scanning Microscope (*Zeiss*) and images obtained at 20X magnification using the Zen Imaging software (*Zeiss*). Analysis of Ki67 positive nuclei was performed using ImageJ software (v1.53a). 10 images per section, at 20X magnification were obtained. Brown (Ki67 positive) and blue (hematoxylin) nuclei were separated using the Color Deconvolution tool with Vectors set to H DAB (Supplemental Diagram 1A). The Threshold function was then used to select nuclei within the corresponding hematoxylin or DAB images, Binary -> Watershed tool was used to separate nuclei where appropriate, and the Measure Particles tool was used to count the number of nuclei within the images. Percent positive nuclei for Ki67 was then calculated, and an average across the 10 images obtained per section.

For analysis of hepatocyte apoptosis, the Click-iT™ Plus TUNEL Assay Kit (*Invitrogen*; Alexa Fluor^TM^ 647) was used following manufacturers protocol. Briefly, 3 µm thick sections were obtained within 24 h of staining, and dewaxed and rehydrated following manufacturers specifications. Sections were permeabilized with proteinase K solution for 15 mins at room temperature, incubated with a TdT reaction solution for 60 mins at 37°C followed by incubation with the Click-iT^TM^ Plus reaction solution 30 mins at 37°C. A positive control was included by incubating a liver section with DNase to induce DNA strand breaks prior to the TdT incubation, and a negative control was included that omitted incubation with the TdT reaction solution. Nuclei were counter stained with DAPI, and staining was visualized using the Axioscan Scanning Microscope (*Zeiss*) and Zen Imaging software (*Zeiss*). For quantification of TUNEL positive cells, 10 images per section at 40X were obtained, keeping the DAPI and 647 channel images separate. DAPI images were imported into ImageJ software (v1.53a) and converted to 8-bit (Supplemental Diagram 1B). The Threshold function was then used to select nuclei, Binary -> Watershed tool was used to separate nuclei where appropriate, and the Measure Particles tool was used to count the number of nuclei. The number of TUNEL positive cells was counted manually. Percent TUNEL positive nuclei was then calculated, and an average across the 10 images obtained per section.

### Quantitative PCR (qPCR)

Approximately 30 mg of snap frozen liver tissue was homogenized and lysed in RLT-lysis buffer (*Qiagen*) aided by a tissue homogenizer. RNA was extracted using the RNeasy Mini Plus kit (*Qiagen*) following standard manufacturers protocol. A total of 1 µg of RNA was converted to complementary DNA (cDNA) using the High-capacity cDNA Reverse Transcription kit (*Applied Biosystems*) and diluted to 1:100. For qPCR, 2.5 µl of cDNA was mixed with 10 µl of PowerUp SYBR green (*Applied Biosystems*), 1.2 µl of primers at a concentration of 10 nM, and water to make up a total reaction volume of 20 µl. Primers for genes of interest specific to the guinea pig genome were purchased commercially (*Millipore* KiCqStart Sybr Primers) or obtained from previously published literature (11, 12, 25, 26). Stability of reference genes *β-actin*, *Gapdh* and *Rsp20* (25) in fetal liver tissue was determined to be 0.013, 0.018 and 0.028, respectively, using Normfinder (35), and gene expression was normalized to the geometric mean of all three. Reactions were performed using the Quant3 Real-Time PCR System (*Applied Biosystems*), and relative mRNA expression calculated using the comparative CT method with the Design and Analysis Software v2.6.0 (*Applied Biosystems*).

### Western Blots

Fetal liver tissue was homogenized in ice-cold RIPA buffer containing protease and phosphatase inhibitors. Protein concentrations determined using Pierce™ Coomassie Plus Assay Kit (*Thermo Fisher Scientific*) following manufacturer’s protocol. 30 μg of protein was mixed with Bolt® SDS Loading Buffer (*Invitrogen*) and Reducing Agent (*Invitrogen*) and denatured by heating at 95°C for 10 min. The lysates and a pre-stain protein ladder (PageRuler, *Thermo Fisher Scientific*) were then run on a 4-12% Tris-Bis precast gel (*Invitrogen*) following manufacturers protocols and transferred onto nitrocellulose membranes using the Bolt Mini Electrophoresis unit (*Invitrogen*). Membranes were placed into 5% skim-milk in Tris-buffered Saline containing Tween 20 (TBS-T) and incubated overnight at 4°C. Primary antibodies (H2A.X (Ser139): 1:500, *Abcam ab131385*; Deptor: 1:1500, *LSBio C187268*; Total Akt: 1:2000, *Cell Signaling 9272*; Phosphorylated Akt (Ser473): 1:2000, *Cell Signaling 4060*) were applied for 2 h at room temperature, the membranes were then washed 3 times in fresh TBS-T, and then further incubated with a HRP conjugated secondary (*Cell Signaling 7074* and *7076*, 1:2000) for 1 h at room temperature. Protein bands were visualized by chemiluminescence using SuperSignal West Femto Maximum Sensitivity Substrate (*Thermo Fisher Scientific*) on the Chemidoc Imager (*Bio-Rad*) and signal intensity of the protein bands calculated using Image Lab software (version 6.1, *Bio-Rad*), normalized to total protein.

### Serum and Liver Assays

Heparinized fetal blood samples were collected at time of sacrifice by cardiac puncture and plasma obtained by centrifugation. For circulating total and direct bilirubin concentrations, 5 µL of plasma was analyzed using a colorimetric assay kit (*Abcam ab235627*) following manufacturers protocol. For circulating total and free cholesterol concentrations, 5 µL of plasma was analyzed using a colorimetric assay kit (*Abcam ab65390*) following manufacturers protocol. For fetal liver total and free cholesterol content, ∼10 mg of flash-frozen tissue was homogenized using a tissue homogenizer, diluted 1:5 and measured using a colorimetric assay kit (*Abcam ab65390*). Fetal liver (∼100 mg flash-frozen tissue) triglyceride content was measured using a fluorometric assay kit (*Abcam ab65336*) following manufactures protocol. For all assays, samples were run in duplicate, concentrations were calculated from a trendline equation based on standard curve data and optical density corrected by subtracting a blank/background read.

### Statistics

All statistical analyses were performed using SPSS Statistics 29 software. Female and male fetuses were analyzed separately. Distribution assumptions were checked with a Q-Q-Plot. To include the possible unmeasurable correlation of the Dam within the litters, statistical significance was determined using generalized estimating equations with gamma as the distribution and log as the link function. Diet (Control or MNR), treatment (Sham or *hIGF1* nanoparticle) and the interaction between diet and treatment were included as main effects. Gestational age and litter size were included as covariates. Direct injection of the placenta compared to indirect exposure to the *hIGF1* or sham nanoparticles was assessed. In the sham treatment, there was no effect of direct placental injection on any outcomes measured and was therefore removed as a main effect in these groups (26). With *hIGF1* nanoparticle treatment, outcomes in male fetuses whose placentas were directly injected different significantly when compared to indirectly exposure male fetuses and therefore remained as a main effect for this comparison. Statistical significance was considered at P ≤ 0.05. For statistically significant results, a Bonferroni post hoc analysis was performed. Results are reported as estimated marginal means ± 95% confidence interval.

## RESULTS

### Placental *hIGF1* nanoparticle treatment improves fetal growth and is associated with changes to fetal liver weight but not hepatocyte proliferation or apoptosis

Maternal, fetal and placental outcomes have been published previously (26). To summarize, no adverse maternal health outcomes or pregnancy losses were recorded. Average litter size was no different between Control and MNR diets (26); one dam (MNR + *IGF1*) with a litter size of 1 was excluded from the study due to significant physiological differences between monozygotic and polyzygotic pregnancies. No resorptions were recorded. All dams became pregnant, 76% (4 Control and 9 MNR) on the first mating and 24% (2 Control and 2 MNR) on the second mating. There was no difference in the number of female or male fetuses between Control or MNR diet (26). However, it was determined at time of sacrifice that only placentas of male fetuses had received a direct placental injection of *hIGF1* nanoparticle. Repeated placental *hIGF1* nanoparticle treatment resulted in gene expression of plasmid-specific *hIGF1* in all placentas of the litter, with placentas from male fetuses that received the direct injection displaying higher levels of *hIGF1* than female and male littermates that were indirectly exposed (26). Placental *hIGF1* nanoparticle treatment in MNR dams restored near-term fetal weight to sizes comparable to sham treated Control dams for both female and male fetuses, increased fetal blood glucose and decreased fetal blood cortisol (26). We have previously shown using in situ hybridization for plasmid-specific *hIGF1* no off-target nanoparticle expression of *hIGF1* in fetal brain and liver following a single *hIGF1* nanoparticle treatment to the placenta (36). Similarly, analysis using qPCR confirmed no expression (no detectable signal after 40 cycles) of plasmid-specific *hIGF1* in any fetal liver tissue following multiple placental *hIGF1* nanoparticle treatments further reiterating no off-target expression of nanoparticle within the fetus (Supplemental Figure S1).

Fetal liver weight at sacrifice was measured to indirectly assess liver growth as reduced liver weight at the beginning of placental *hIGF1* nanoparticle treatment has been shown previously (37). In female fetuses, liver weight was reduced in sham treated MNR and indirectly exposed MNR + *hIGF1* when compared to Control (Table 1). When liver weight was expressed relative to fetal body weight, sham treated Control and sham treated MNR female fetuses were comparable (26). However, because fetal weight was higher in the indirectly exposed MNR + *hIGF1* females, liver weight as a percentage of fetal weight was decreased when compared to sham treated MNR (26). In male fetuses, liver weight was comparable between sham treated Control, sham treated MNR and direct injected MNR + *hIGF1* but increased in indirectly exposed MNR + *hIGF1* (Table 1). Consequently, when compared to sham treated Controls, liver weight as a percentage of fetal weight was increased in the sham treated MNR male fetuses because fetal weight was lower, and increased in the indirectly exposed MNR + *hIGF1* because liver weight was higher (26).

**Table 1.**
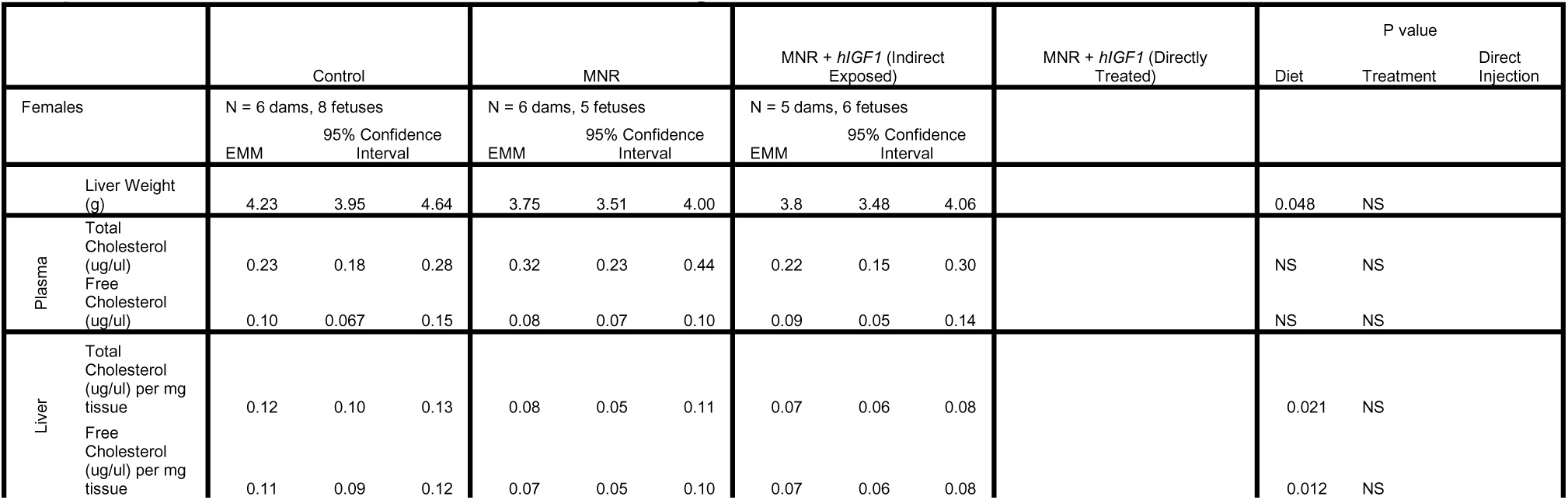
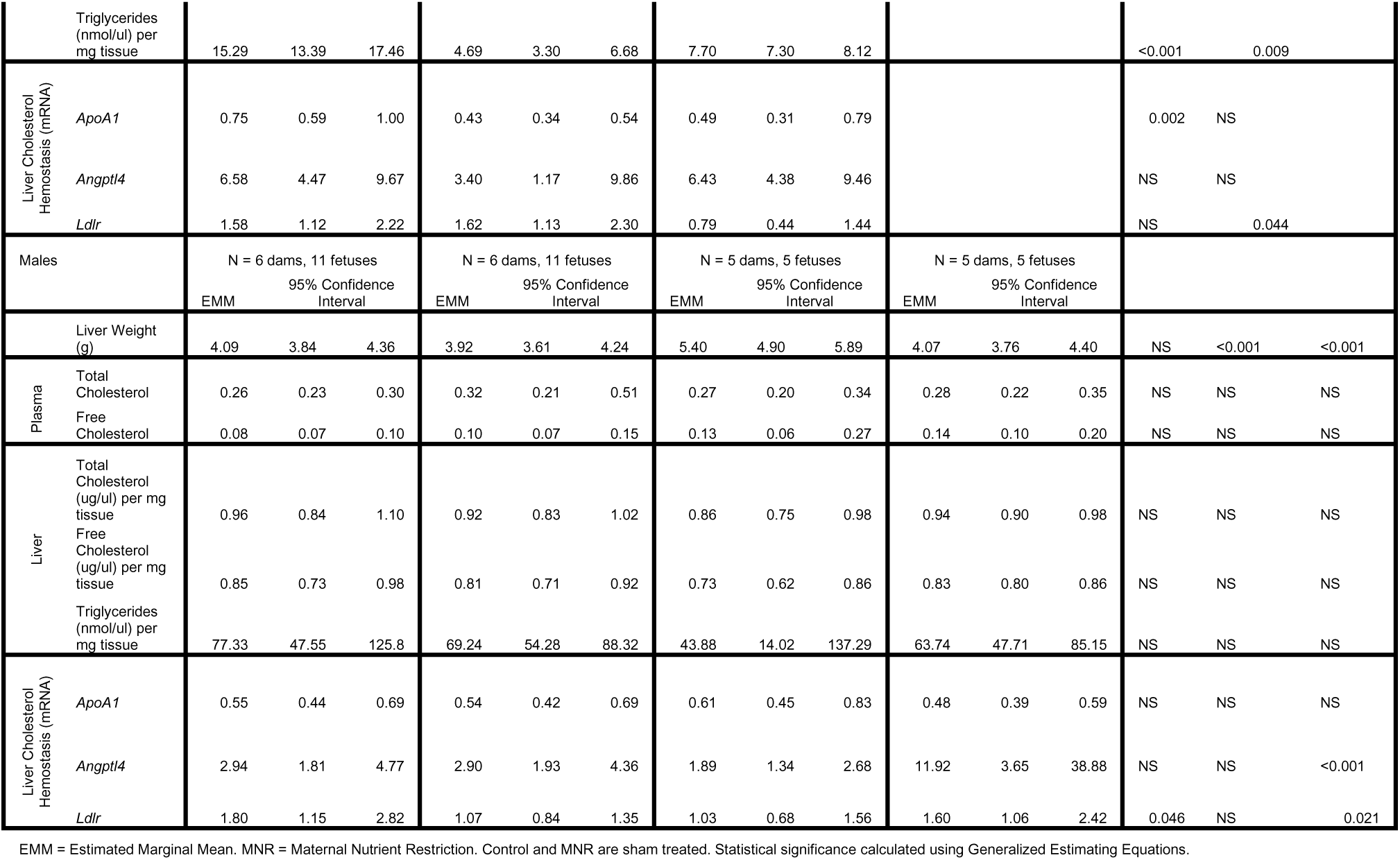
Fetal liver weight, plasma and liver cholesterol and triglycerides and liver expression of cholesterol homeostasis genes.

Assessment of liver portal triad morphology and lipid accumulation in female and male liver tissue demonstrated very little differences with MNR and placental *hIGF1* nanoparticle treatment (Supplemental Figure S2). Collagen deposition surrounding the portal triads was similar between the different groups in both female and male fetuses (Supplemental Figure S3). Similarly, the number of proliferative hepatocytes did not change with either MNR or placental *hIGF1* nanoparticle treatment (Supplemental Figure S4). Finally, the number of TUNEL positive cells in liver tissue, and liver protein expression of H2A.X (pSer139), a biomarker of double-stranded DNA damage, was similar between sham treated Control, sham treated MNR and placental *hIGF1* nanoparticle treated MNR female and male fetuses (Supplemental Figure S5).

### Placental *hIGF1* nanoparticle treatment indirectly influences FGR-associated changes in lipid homeostasis and cholesterol metabolism-related genes in the fetal liver dependent on fetal sex

MNR reduced maternal circulating total cholesterol in both the sham treated MNR and placental *hIGF1* nanoparticle treated MNR dams when compared to sham treated Control (Estimated Mean (µg/µl), 95% CI: Control = 2.55, 2.23-2.86 vs. MNR = 1.86, 1.54-2.18 vs. MNR + *hIGF1* = 1.86, 1.52-2.21, P=0.003). In the fetus, circulating levels of total and free cholesterol were similar across groups in both female and male fetuses (Table 1). Per mg of liver tissue, MNR reduced liver total and free cholesterol and triglycerides in female fetuses when compared to sham treated Control (-34%, -37% and -69%, respectively; Table 1). Liver total and free cholesterol and triglycerides were also lower in MNR female fetuses whose placentas were indirectly exposed to the placental *hIGF1* nanoparticle treatment when compared to sham treated Control (-37%, -35% and -50%, respectively; Table 1), however, triglycerides were increased compared to sham treated MNR (+64%; Table 1). Liver cholesterol metabolism-related gene expression of *ApoA1* was decreased in the sham treated MNR and indirectly exposed MNR + *hIGF1* female fetuses compared to sham treated Control (-43% and -35%, respectively; Table 1), *Angptl4* was similar across all groups, and *Ldlr* was decreased in the indirectly exposed MNR + *hIGF1* female fetal liver compared to sham treated Control and sham treated MNR (-50% and -51%, respectively; Table 1). In male fetuses, per mg of tissue, liver total and free cholesterol and triglycerides were similar across all groups (Table 1). Liver cholesterol homeostasis gene expression of *ApoA1* was similar across all groups, whilst expression of *Angptl4* and *Ldlr* was increased in the livers of males whose placenta were directly treated with placental *hIGF1* nanoparticle compared to those whose placentas were indirectly exposed to the *hIGF1* nanoparticle (+530% and +56%, respectively; Table 1).

### FGR-associated changes to fetal liver gluconeogenesis and glucose metabolism-related signaling pathway genes are prevented with placental *hIGF1* nanoparticle treatment dependent on fetal sex

Endogenous glucose production is increased with FGR to compensate for reduced placental glucose supply (17), hence hepatic expression of glucose-metabolism signaling pathway genes and proteins was assessed. In female fetuses, liver protein expression of the mTOR inhibitor Deptor, Akt and Akt phosphorylation status was similar between sham treated Control, sham treated MNR and indirectly exposed MNR + *hIGF1* (Figure 1A-C). In male fetuses, liver protein expression of Deptor was decreased in directly treated MNR + *hIGF1* fetuses compared to all other groups (-67%, -57% and -47%, respectively; Figure 1D). Protein expression of Akt and Akt phosphorylation status was similar between the groups (Figure 1E and 1F, respectively). Representative western blot images are presented in Supplemental Figure S6. In addition to changes in liver PI3K/Akt signaling proteins, in liver tissue from female fetuses, gene expression of gluconeogenesis enzymes *G6pc* and *Pck2*, and *IgfBP3* was increased in sham treated MNR compared to sham treated Control (+70%, +43% and +76%, respectively; Figure 2A-2C), but no different from sham treated Control in the MNR female fetuses indirectly exposed to the placental *hIGF1* nanoparticle. Gene expression of *Igf1* was decreased in the sham treated MNR and indirectly exposed MNR + *hIGF1* female fetuses when compared to sham treated Control (-26% and -55%, respectively; Figure 2D), whilst expression of *IgfBP1* was increased in the indirectly exposed MNR + *hIGF1* female fetuses compared to sham treated MNR (+22%, Figure 2E) and similar to sham treated Control. Female fetal liver gene expression of *Slc2A1* was similar across the groups (Figure 2F). In liver tissue from male fetuses, gene expression of *G6pc*, *Pck2* and *IgfBP3* was decreased with placental *hIGF1* nanoparticle treatment in directly treated male fetuses when compared to sham treated MNR (-30%, -43% and -50%, respectively; Figure 2G-I). Gene expression of *Igf1* was decreased in male fetal liver with both indirect exposure to *hIGF1* nanoparticle and direct injection of *hIGF1* nanoparticle when compared to sham treated Control (-32% and -33%, respectively; Figure 2J). The gene expression of *IgfBP1* was similar across groups (Figure 2K), whilst gene expression of *Slc2A1* was increased in sham treated MNR and MNR + *hIGF1* male fetuses when compared to sham treated Control (+109%, +117% and +251%, respectively; Figure 2L). In both females and males, mRNA expression of *Pck1*, *GcgR*, *Igf1R* and *Igf2* were similar across groups (Supplemental Table S1).

**Figure 1.**
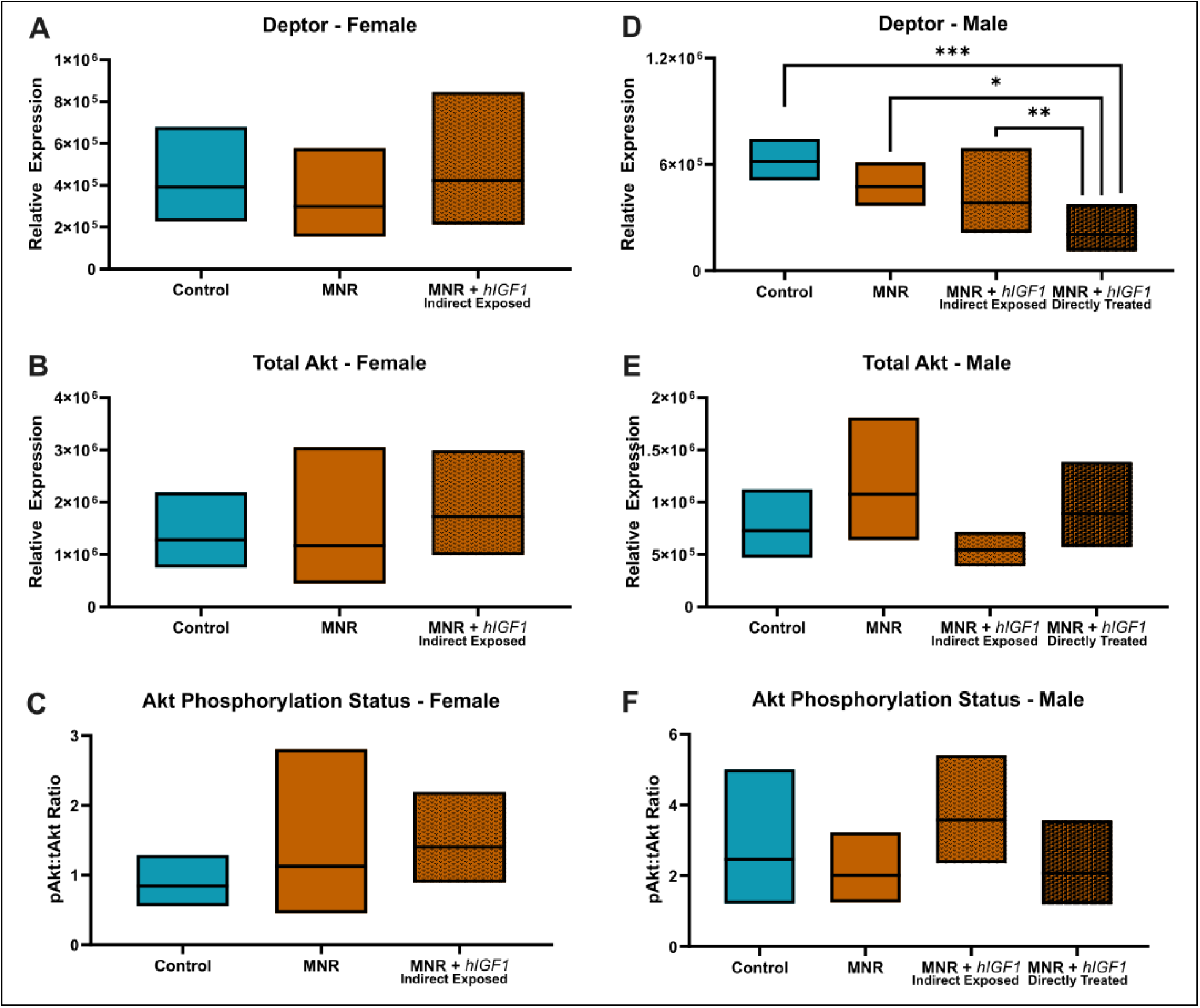
Effects of maternal nutrient restriction (MNR) and repeated placental *hIGF1* nanoparticle treatment (MNR + *hIGF1*) on fetal liver protein expression of PI3K/AKT signaling factors. **A.** In female fetuses, protein expression of the mTOR inhibitor Deptor was similar across the groups. **B.** In female fetuses, protein expression of Total Akt was similar across the groups. **C**. In female fetuses, Akt phosphorylation status was similar across the groups. **D.** In male fetuses, protein expression of the mTOR inhibitor Deptor was decreased in directly treated MNR + *hIGF1* fetuses compared to sham treated Control, sham treated MNR and indirectly exposed MNR + *hIGF1* fetuses **E.** In male fetuses, protein expression of Total Akt was similar across the groups. **F.** In male fetuses, Akt phosphorylation status was similar across the groups. Control: n = 6 dams (8 female and 11 male fetuses), MNR: n = 6 dams (5 female and 11 male fetuses), MNR + *hIGF1*: n = 5 dams (6 female and 10 male fetuses). Data are estimated marginal means ± 95% confidence interval. *P≤0.05; **P≤0.01. ***P≤0.001

**Figure 2.**
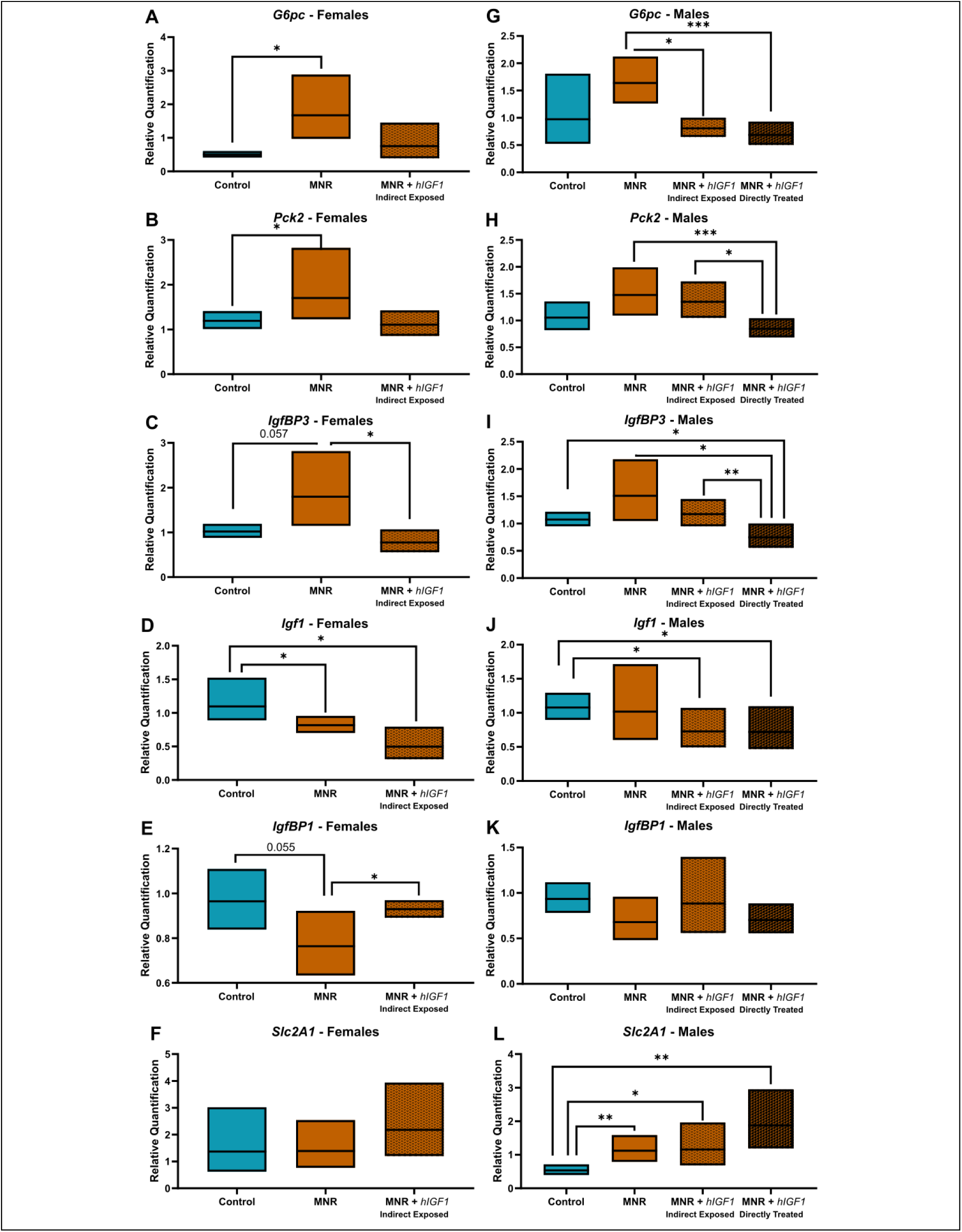
Effects of maternal nutrient restriction (MNR) and repeated placental *hIGF1* nanoparticle treatment (MNR + *hIGF1*) on fetal liver expression of gluconeogenesis and glucose metabolism-related signaling pathway genes. **A.** In female fetuses, liver mRNA expression of *Glucose-6-Phosphatase* (*G6pc*) was increased in sham treated MNR fetuses when compared to Control, but similar in MNR + *hIGF1* fetuses when compared to Control. **B.** In female fetuses, liver mRNA expression of *Phosphoenolpyruvate Carboxykinase 2* (*Pck2*) was increased in sham treated MNR fetuses when compared to Control, but similar in MNR + *hIGF1* fetuses when compared to Control. **C**. In female fetuses, liver mRNA expression of *Insulin-like Growth Factor Binding Protein 3* (*IgfBP3*) was decreased in MNR + *hIGF1* fetuses when compared to sham treated MNR **D.** In female fetuses, liver mRNA expression of *Insulin-like Growth Factor 1* (*Igf1*) was decreased in sham treated MNR and MNR + *hIGF1* fetuses when compared to Control **E.** In female fetuses, liver mRNA expression of *IgfBP1* was increased in MNR + *hIGF1* fetuses when compared to sham treated MNR **F.** In female fetuses, liver mRNA expression of *Glucose Transporter 1* (*Slc2A1*) was similar across the groups. **G.** In male fetuses, liver mRNA expression of *G6pc* was decreased in both indirectly exposed and directly treated MNR + *hIGF1* treated fetuses compared to sham treated MNR fetuses. **H.** In male fetuses, liver mRNA expression of *Pck2* was decreased in directly treated MNR + *hIGF1* treated fetuses compared to sham treated MNR and indirectly exposed MNR + *hIGF1* fetuses. **I**. In male fetuses, liver mRNA expression of *IgfBP3* was decreased in directly treated MNR + *hIGF1* treated fetuses compared to Control, sham treated MNR and indirectly exposed MNR + *hIGF1* fetuses. **J.** In male fetuses, liver mRNA expression of *Igf1* was decreased in indirectly exposed and directly treated MNR + *hIGF1* treated fetuses compared to Control. **K.** In male fetuses, liver mRNA expression of *IgfBP1* was similar across the groups. **L.** In male fetuses, liver mRNA expression of *Slc2A1* was increased in sham treated MNR, indirectly exposed and directly treated MNR + *hIGF1* fetuses when compared to control. Control: n = 6 dams (8 female and 11 male fetuses), MNR: n = 6 dams (5 female and 11 male fetuses), MNR + *hIGF1*: n = 5 dams (6 female and 10 male fetuses). Data are estimated marginal means ± 95% confidence interval. *P≤0.05; **P≤0.01. ***P≤0.001

### FGR is associated with increased liver gene expression of pro-fibrosis markers, but prevented with placental *hIGF1* nanoparticle treatment

Increased transcriptional activation of hepatic fibrosis-related genes an important predictor of liver dysfunction (38–41) which is further mediated by ‘second hits’ such increased inflammation (42). Gene expression of pro-fibrosis markers *Hif1a* and *Mmp2* were decreased with placental *hIGF1* nanoparticle treatment in livers of indirectly exposed females (-40% and -39%, respectively) and directly treated males (-48% and -40%, respectively) when compared to sham treated MNR, and comparable to sham treated Control (Figure 3A-E). Expression of *Tgfb* in directly treated males was also decreased compared to sham treated MNR and comparable to levels in sham treated Control (-40%; Figure 3F). mRNA expression of other pro-fibrosis markers *Ctgf*, *Timp1* and *Timp2*, was similar between groups in both female and male fetal livers (Supplemental Table S1).

**Figure 3.**
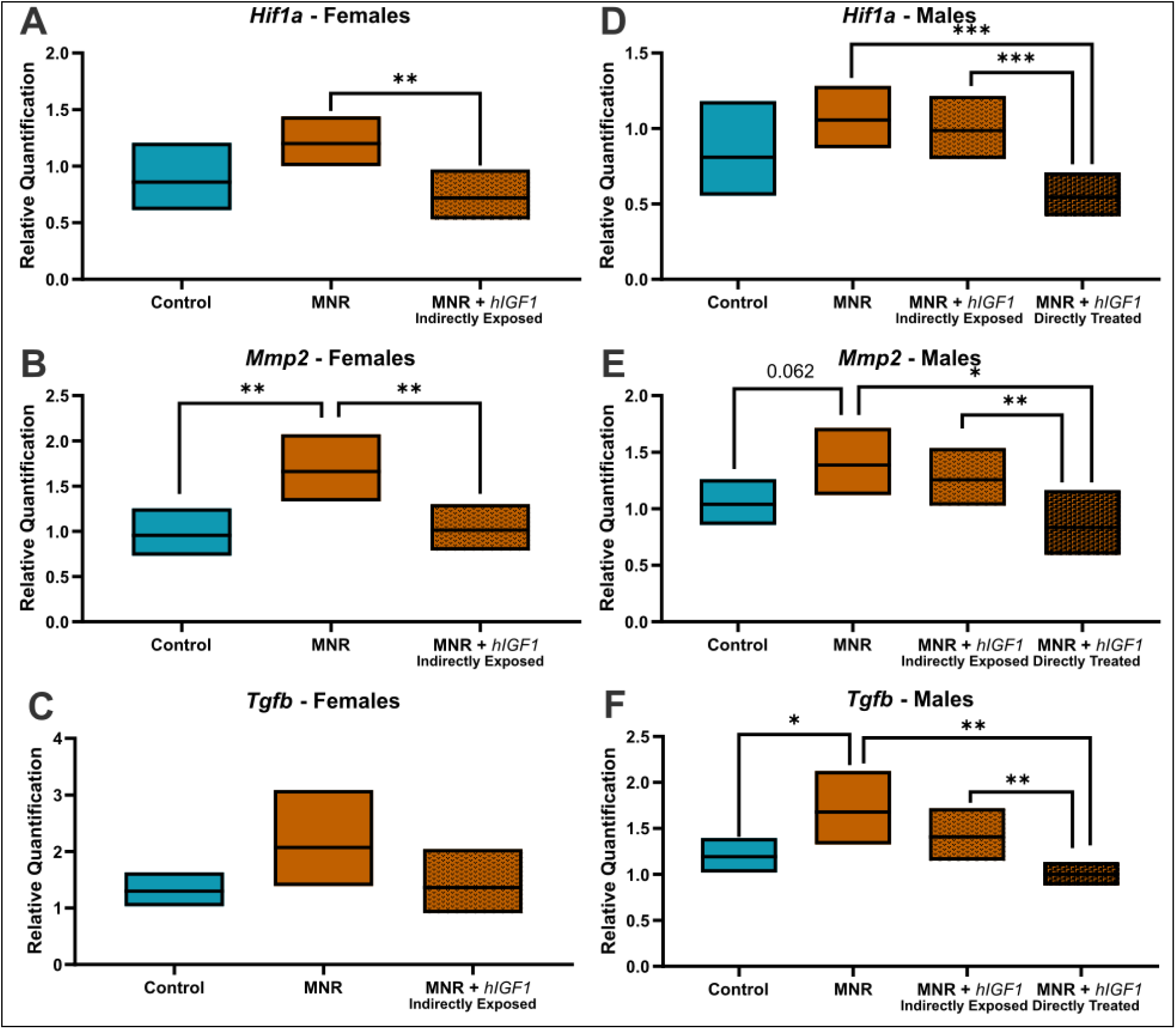
Effects of maternal nutrient restriction (MNR) and repeated placental *hIGF1* nanoparticle treatment (MNR + *hIGF1*) on fetal liver gene expression of pro-fibrotic markers. **A.** In female fetuses, mRNA expression of *Hypoxia Inducible Factor 1a* (*Hif1a*) was decreased in MNR + *hIGF1* livers compared to sham treated MNR. **B.** In female fetuses, mRNA expression of *Matrix Metalloprotein 2* (*Mmp2*) was increased in sham treated MNR compared to Control and decreased in MNR + *hIGF1* livers compared to sham treated MNR. **C**. In female fetuses, whilst there was no statistically significant changes to the expression of *Transforming Growth Factor beta* (*Tgfb*), a similar trend in expression levels was seen as *Mmp2* and *Hif1a*. **D.** In male fetuses, liver expression of *Hif1a* was decreased in directly treated MNR + *hIGF1* fetuses compared to sham treated MNR and indirectly exposed MNR + *hIGF1*. **E.** In male fetuses, liver expression of *Mmp2* was decreased in directly treated MNR + *hIGF1* fetuses compared to sham treated MNR and indirectly exposed MNR + *hIGF1*. **F.** In male fetuses, liver expression of *Tgfb* was increased in sham treated MNR compared to Control and decreased in directly treated MNR + *hIGF1* fetuses compared to sham treated MNR and indirectly exposed MNR + *hIGF1*. Control: n = 6 dams (8 female and 11 male fetuses), MNR: n = 6 dams (5 female and 11 male fetuses), MNR + *hIGF1*: n = 5 dams (6 female and 10 male fetuses). Data are estimated marginal means ± 95% confidence interval. *P≤0.05; **P≤0.01. ***P≤0.001

Increased immune cell infiltration was not observed around the portal triads with either MNR or placental *hIGF1* nanoparticle treatment in either female or male fetuses (Supplemental Figure S7). Similarly, liver gene expression of inflammatory markers *Tnfa*, *Il6* and *Il6 Receptor* (*Il6R*) was not different across the groups for both females and males (Supplemental Table S1). However, expression of *Il1b* was decreased with MNR in sham treated and indirectly exposed *hIGF1* nanoparticle treated females compared to sham treated Control (-43% and -65%, respectively; Supplemental Table S1).

### Placental *hIGF1* nanoparticle treatment increases production of direct bilirubin in FGR fetuses

Plasma total and direct bilirubin were measured to indirectly assess functional metabolic changes in the fetal liver (43). In female fetuses, whilst there was no difference in circulating total bilirubin concentrations (Figure 4A), direct bilirubin was decreased between sham treated MNR and sham treated Control (-15%; Figure 4B). Placental *hIGF1* nanoparticle treatment increased both total and direct bilirubin concentrations when compared to sham treated MNR (+46% and +31%, respectively) and was comparable to sham treated Control. In male fetuses, circulating levels of total and direct bilirubin appeared similar between sham treated Control and sham treated MNR (Figure 4C and 4D). In male fetuses whose placentas were *hIGF1* nanoparticle treated, circulating levels of total and direct bilirubin did not reach a statistically significant increase (P≤0.05). However, Wald’s chi-square: total bilirubin = 2.70 (95% Wald CI = -0.65 to 0.057) and direct bilirubin = 4.57 (95% Wald CI = -1.023 to -0.44) indicated a potential association between placental *hIGF1* nanoparticle treatment and circulating bilirubin concentrations.

**Figure 4.**
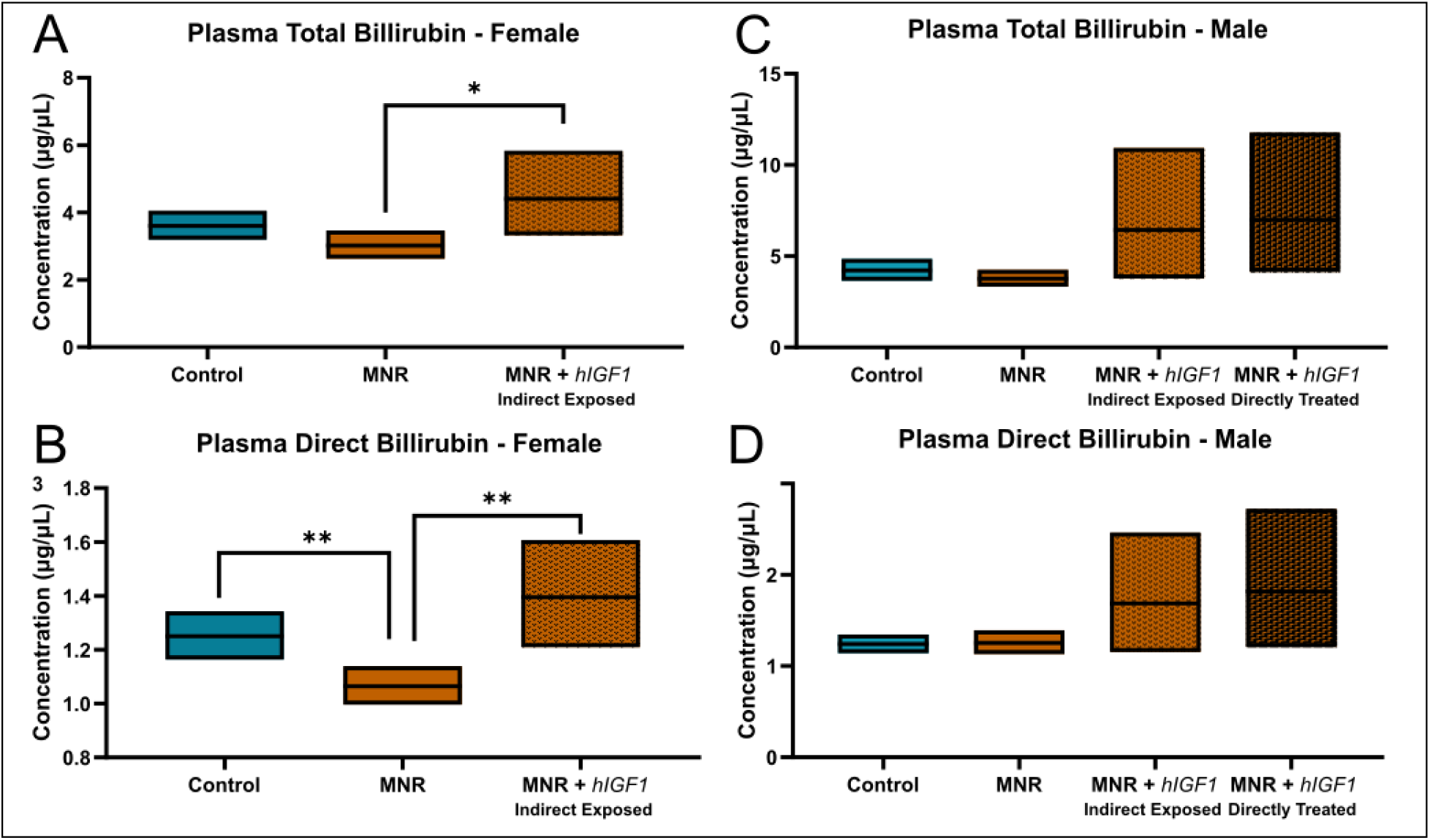
Effects of maternal nutrient restriction (MNR) and repeated *hIGF1* nanoparticle gene therapy (MNR + *hIGF1*) on fetal circulating total and direct bilirubin. **A.** In female fetuses there was no difference in circulating total bilirubin concentrations between sham treated MNR and Control but *hIGF1* nanoparticle treatment increased total when compared to sham treated MNR. **B.** In female fetuses, direct bilirubin was decreased with sham treated MNR when compared to Control but was increased in MNR + *hIGF1* fetuses when compared to MNR and was comparable to Control. **C & D.** In male fetuses, circulating levels of total bilirubin appeared similar between sham treated Control and MNR. However, in male fetuses whose placentas were *hIGF1* nanoparticle treated, whilst circulating levels of total and direct bilirubin did not reach a statistically significant increase, Wald’s chi-square indicated a potential association between *hIGF1* nanoparticle treatment and increased circulating bilirubin levels in male fetuses that may be more clearly determined with a larger sample size. Control: n = 6 dams (8 female and 11 male fetuses), MNR: n = 6 dams (5 female and 11 male fetuses), MNR + *hIGF1*: n = 5 dams (6 female and 10 male fetuses). Data are estimated marginal means ± 95% confidence interval. *P≤0.05; **P≤0.01.

## DISCUSSION

Using the well characterized guinea pig maternal nutrient restriction (MNR) model of FGR and building on previous placental *hIGF1* nanoparticle feasibility studies (25, 26, 37, 44), our aim was to determine the impact of a placenta-specific *hIGF1* nanoparticle gene therapy and the associated improved fetal weight on fetal liver growth and activity and expression of hepatic lipid and glucose metabolism-related signaling pathways gene and proteins in the near-term fetal liver. We demonstrated that the placental *hIGF1* nanoparticle treatment resulted in changes to liver size, cholesterol homeostasis and glucose/nutrient metabolism-related genes in the growth restricted fetus that might confer protection against increased susceptibility to aberrant liver physiology in later-life. Impact on fetal liver physiology was due to enhanced placenta efficiency as neither the nanoparticle nor plasmid crossed the placenta to fetal circulation. The changes in liver gene and protein expression, and indirect measures of hepatic function occurred in a sexually dimorphic manner offering further insight into the unique responses of female and male fetuses to adverse in utero environments. Interestingly, changes to pro-fibrosis markers and nutrient sensing genes in male fetuses was highly influenced by level of placental *hIGF1* expression shown previously (26), highlighting a potential “dose-response” relationship whereby higher placental *hIGF1* expression resulted in more robust changes to liver gene expression. Overall, the findings illuminate the transformative potential of the placental *hIGF1* nanoparticle gene therapy in ameliorating the impacts of placenta insufficiency on fetal liver development, suggesting a promising therapeutic avenue for addressing FGR-related metabolic programming challenges.

Throughout life, the liver is central to coordinating many metabolic pathways including maintaining appropriate glucose levels (45), and in utero programming of impaired glucose tolerance and insulin resistance is well characterized from experimental studies (18, 46–49). These studies highlight the vulnerability of fetal hepatic development particularly due to undernutrition because of failure of the placenta to appropriately transport nutrients and survival adaptations fetus undergoes. In the present study, liver weight was reduced in the growth restricted female fetus and not corrected, despite increased overall fetal weight, with placental *hIGF1* nanoparticle treatment. Reduced liver growth was not a consequence of reduced proliferation or increased apoptosis but associated with decreased expression of *Igf1* and no changes to expression of glucose transporter *Slc2A1*. Prior investigations on the impact of placental insufficiency on fetal liver growth demonstrate under hypoglycemic conditions where liver growth is maintained, upregulation of *Slc2A1* expression as a possible compensatory mechanism to increased glucose uptake in the liver (16). Indeed, we have shown reduced circulating glucose in the FGR female and male fetus in our model (26) and our results in the female fetus support the hypothesis that in the absence of increased *Slc2A1* expression and compensatory glucose uptake, expression of *Igf1* is reduced and possibly contributes to reduced liver growth. On the other hand, in FGR male fetuses, liver weight was increased relative to fetal weight and associated with increased expression of *Slc2A1* whilst *Igf1* expression was variable. Overall, these results highlight fundamental mechanistic differences in liver development between female and male fetuses that may confer with observations that male offspring are at higher risk for developing glucose intolerance and insulin resistance in later life (18).

Disruptions to lipid homeostasis contribute to liver pathophysiology and conditions such as dyslipidemia and non-alcoholic fatty liver disease. In particular, issues with cholesterol homeostasis result in cholesterol accumulation in the liver, especially of the free unesterified cholesterol, resulting in a lipotoxic state (50). Circulating total and free cholesterol concentrations were similar between normal growing and growth restricted near-term fetuses, a finding similar to what has been shown in growth restricted guinea pig offspring at young adulthood (51). However, liver total and free cholesterol and triglycerides were reduced in the growth restricted female fetuses. Reduced lipid concentrations in the liver were associated with reduced expression of *ApoA1*, an apolipoprotein of high-density lipoprotein (HDL) which regulates lipid transport through the reverse cholesterol transport pathway (52). Together, these results suggest a possible mechanism for preserving circulating lipid concentrations to ensure adequate lipid availability to support the growth of peripheral tissues. Liver cholesterol, triglycerides and *ApoA1* expression remained reduced in the female fetuses whose placentas were indirectly exposed to the *hIGF1* nanoparticle gene therapy, and possibly driven by reduced lipid availability from maternal circulation. We have previously demonstrated upregulation of glucose and amino acid transport mechanisms in the placenta following *hIGF1* gene therapy treatment (25). However, the effect of placental *hIGF1* treatment on lipid transport in the placenta has not thoroughly been investigated, and beyond the scope of the current manuscript. Interestingly, the contrast in liver lipid profiles and cholesterol homeostasis gene expression observed between male and female fetuses indicates sexual dimorphism in fetal liver responses to in utero challenges. Male fetuses showed similar liver lipid levels across all groups, and differential gene expression predominantly in response to direct placental *hIGF1* nanoparticle treatment. These findings parallel previous studies showing that only growth restricted male guinea pigs exhibit alterations in cholesterol metabolism-related signaling pathway genes in the liver in young adulthood, and not females (12). Building on from this previous study we show reduced liver gene expression of *Ldlr* in the growth restricted males at young adulthood is programmed prenatally, and in the absence of elevated liver cholesterol concentrations. Additionally, liver expression of *Ldlr* was increased in the growth restricted males whose placentas received a direct placental *hIGF1* nanoparticle treatment and comparable to normal growing fetuses, further reiterating the potential for this therapy in mitigating against disrupted lipid homeostasis.

One of the prevailing hypotheses as to why people born growth restricted are at increased risk of glucose intolerance and insulin resistance is because the liver is programmed during fetal development to increase endogenous glucose production to support appropriate fetal growth (17, 53). The programmed increased endogenous glucose production then persists after birth and is mis-matched with the postnatal environment, resulting in issues with glucose metabolism. Here, we show changes to protein expression of PI3K/AKT signaling proteins because of placental treatment with the *hIGF1* nanoparticle gene therapy in a sex-dependent manner. The PI3K/AKT signaling pathway is one of the major pathways that regulates glucose metabolism and gluconeogenesis in the liver (54). Changes to PI3K/AKT signaling were associated with reduced gene expression of gluconeogenesis enzymes *G6pc* and *Pck1* in fetuses whose placentas were treated with *hIGF1* nanoparticle compared to sham-treated growth restricted fetuses, and more similar levels of expression to normal Control fetuses. Both G6PC and PCK2 are rate limiting enzymes that contribute to the formation of glucose from pyruvate in the liver (55). Increased hepatic gluconeogenesis is associated with hyperglycemia in patients with type 2 diabetes (56) as well as hepatic insulin resistance (57), and animal models confirm increased expression in the liver because of FGR (17, 46). In addition to glucose, cortisol has been shown to regulate expression of gluconeogenesis markers in the liver (58). We have previously shown the ability to reduce circulating fetal cortisol with placental *hIGF1* nanoparticle treatment, thereby dampening the fetal stress response (26). Hence, our data shows the likely potential for the *hIGF1* nanoparticle gene therapy to mitigate increased risk of glucose intolerance and insulin resistance in later life by preventing programmed increased endogenous glucose production.

Liver fibrosis characterized by an excessive deposition and reorganization of extracellular matrix (ECM), is also an important predictor of adverse long-term outcomes in the liver (38–41). Increased collagen deposition was not observed within the livers of both female and male fetuses, likely reflective of the developmental stage as excessive ECM deposition from birth is likely to confer very poor survival. However, liver expression analysis of pro-fibrotic markers *Hif1a*, *Mmp2* and *Tgfb* indicated a potential relationship between growth restriction and increased expression, similar to what has been shown in the livers of growth restricted offspring in young adulthood (11). TGFB1 is known to promote liver fibrosis and upregulate MMP2 expression which further accelerates hepatic fibrosis and disease progression (59, 60). In young adult offspring fed a normal diet post-partum who were born growth restricted, mRNA expression of *Tgfb*, *Ctgf*, and *Mmp2* was markedly increased compared to offspring born at normal weight. These findings together with ours provide further evidence that an adverse in utero environment which results in FGR activates profibrotic pathways in the fetal liver which are sustained into adulthood. Interestingly, liver expression of *Hif1a*, *Mmp2* and *Tgfb* was only observed in the male fetuses whose placentas were directly injected with the placental *hIGF1* nanoparticle, and not male fetuses whose placentas were indirectly exposed. We have previously shown that placenta expression of plasmid-specific *hIGF1* was lower in indirectly exposed placentas compared to those directly injected (26), and together with the results presented here suggest a potential “dose” response in the fetus that is related to placental *hIGF1* expression. Despite also only being indirectly exposed to *hIGF1* nanoparticle and having lower placental expression of plasmid-specific *hIGF1* (26), changes in liver gene expression of pro-fibrotic markers in females was similar to directly treated males, suggesting that females may potential respond better to therapeutic interventions than males.

The influence of fetal sex on pregnancy outcomes is well established, with male fetuses at higher risk of poorer outcomes then female fetuses (61). Hence, it is unsurprising that our results suggest the potential for female fetuses to respond better to therapeutic interventions. However, one limitation to further exploring this hypothesis in the present study is the inability to study the impact of the *hIGF1* nanoparticle gene therapy in female fetuses whose placentas were directly injected. As most human pregnancies are singleton with one placenta, one fetus in the litter was chosen to receive a direct sham or *hIGF1* nanoparticle gene therapy injection to study effects of the full direct treatment. The remaining fetuses in the litter were used to investigate possible circulating indirect exposure. However, the sex of the fetus which receives the direct injection cannot be controlled nor reliably determined at mid-pregnancy via ultrasound. Further studies in non-human primates are being performed to address translational aspects of the *hIGF1* nanoparticle gene therapy including dose, which are difficult to ascertain in litter bearing species (62). Another limitation to this study was the inability to directly measure liver metabolism, glucose tolerance or insulin resistance because of the developmental stage. Circulating bilirubin can be used as an indirect measure of liver function, as unconjugated bilirubin is metabolized in the liver to direct (conjugated) bilirubin which can then be excreted in urine (63). Typically, high unconjugated bilirubin is associated with adverse health outcomes such as hepatitis and jaundice, however, low serum bilirubin is directly correlated with pathological conditions including diabetes mellitus, metabolic syndrome, and cardiovascular diseases (64). In our study, we show increased circulating direct bilirubin in growth restricted female fetuses whose placentas were indirectly exposed to the *hIGF1* nanoparticle treatment, and a potentially similar impact in male fetuses, indicating the possibility that liver function is enhanced. Further studies are focusing on assessing liver function in offspring, including determining the impact of preventing growth restriction with the *hIGF1* nanoparticle gene therapy on glucose tolerance and lipid metabolism in adulthood.

In conclusion, here we investigated the effects of a placental-specific *hIGF1* nanoparticle treatment on fetal liver development, focusing on gene and protein expression of factors regulating hepatic lipid and glucose homeostasis. Our findings underscore the beneficial impact of the placental *hIGF1* nanoparticle treatment and associated improved placenta efficiency on liver growth, cholesterol homeostasis, and nutrient sensing, potentially offering protection against later-life liver pathologies. Interestingly, these alterations occurred in a sexually dimorphic manner, revealing distinct responses between female and male fetuses to in utero adversities. The differences in hepatic development and metabolic programming between fetal sex emphasize the need for sex-specific approaches in understanding and treating FGR-induced metabolic disturbances. The research also illuminates a possible “dose-response” relationship based on placental plasmid-specific *hIGF1* expression, which significantly influenced liver gene expression changes. Overall, given the complex interplay of factors influencing fetal development as well as the translational prospect of mitigating FGR-induced metabolic derangements, our work opens avenues for future research, especially regarding the optimal application of the *hIGF1* nanoparticle treatment in further pre-clinical and human clinical scenarios.

## Acknowledgments

We would like to thank Drs Craig Duvall and Mukesh Gupta for providing the co-polymer, veterinary and technical staff at the University of Florida Animal Care Services, and Dr. Khanh Huynh, Aditya Mahadevan, Dr Jason Puglise, and Dr Erica Smith for their assistance with animal necropsies.

## Author Contributions

BND: Methodology, Investigation, Validation, Resources, Writing – Review & Editing. AW: Investigation, Validation. TRHR: Conceptualization, Methodology, Supervision, Writing – Review & Editing. HNJ: Conceptualization, Resources, Supervision, Funding Acquisition, Writing – Review & Editing. RLW: Conceptualization, Methodology, Validation, Investigation, Formal Analysis, Writing - Original Draft, Project Administration, Funding Acquisition

## Ethics approval

Animal care and usage was approved by the University of Florida Intuitional Animal Care and Usage Committee (Protocol #202011236).

