## Supplemental Material for "Placental *hIGF1* nanoparticle gene therapy in guinea pigs ameliorates fetal growth restriction-associated changes in hepatic lipid and glucose metabolism-related signaling pathways in a fetal sex-specific manner"

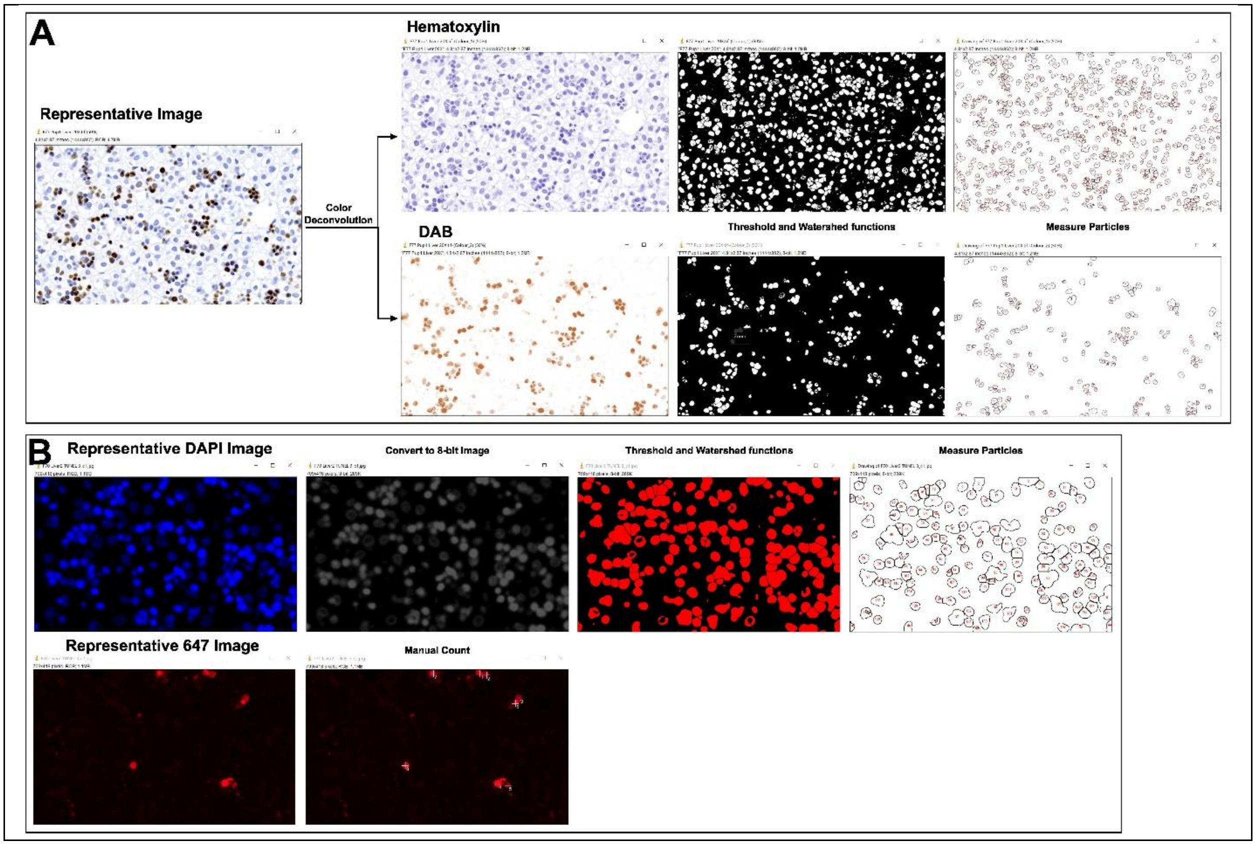


**Supplemental Diagram 1. Representative schematic of immunohistochemistry analysis using ImageJ Software. A.** Brown (Ki67 positive) and blue (hematoxylin) nuclei were separated using the Color Deconvolution tool with Vectors set to H DAB. The Threshold function was then used to select nuclei within the corresponding hematoxylin or DAB images, Binary -> Watershed tool was used to separated nuclei where appropriate, and the Measure Particles tool was used to count the number of nuclei within the images. **B.** DAPI images were imported into ImageJ software and converted to 8-bit. The Threshold function was then used to select nuclei, Binary -> Watershed tool was used to separated nuclei where appropriate, and the Measure Particles tool was used to count the number of nuclei. The number of TUNEL positive cells was counted manually.

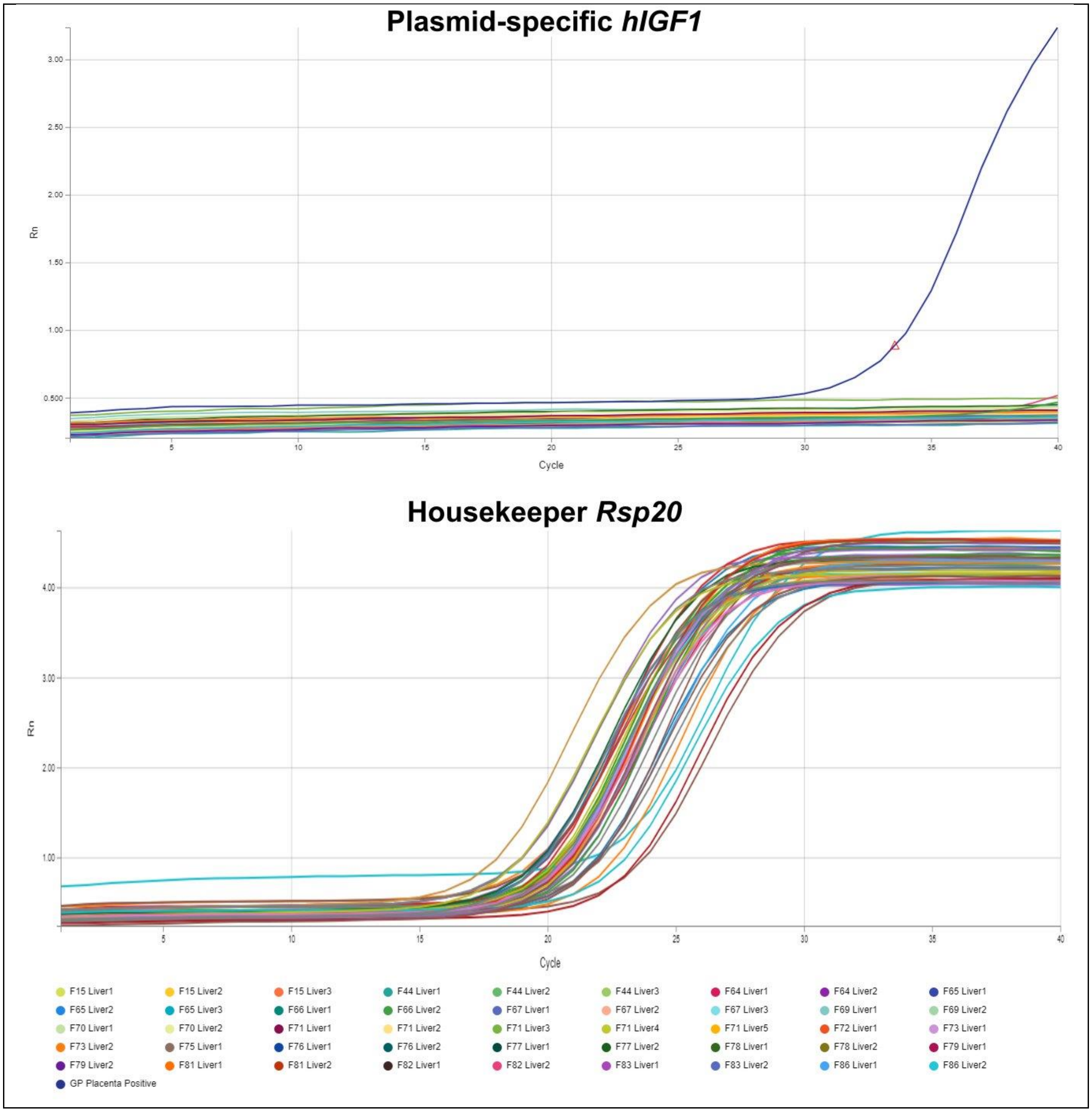


**Supplemental Figure S1. qPCR expression of plasmid-specific human *IGF1* (*hIGF1*) in fetal liver.** There was no expression (no detectable fluorescence after 40 cycles) of plasmid-specific *hIGF1* in any fetal liver tissue following multiple placental *hIGF1* nanoparticle gene therapy treatments. Guinea pig placenta treated with *hIGF1* nanoparticle gene therapy was included as a positive control for *hIGF1*. qPCR for housekeeper gene *Rsp20* confirmed presence of cDNA in all samples.

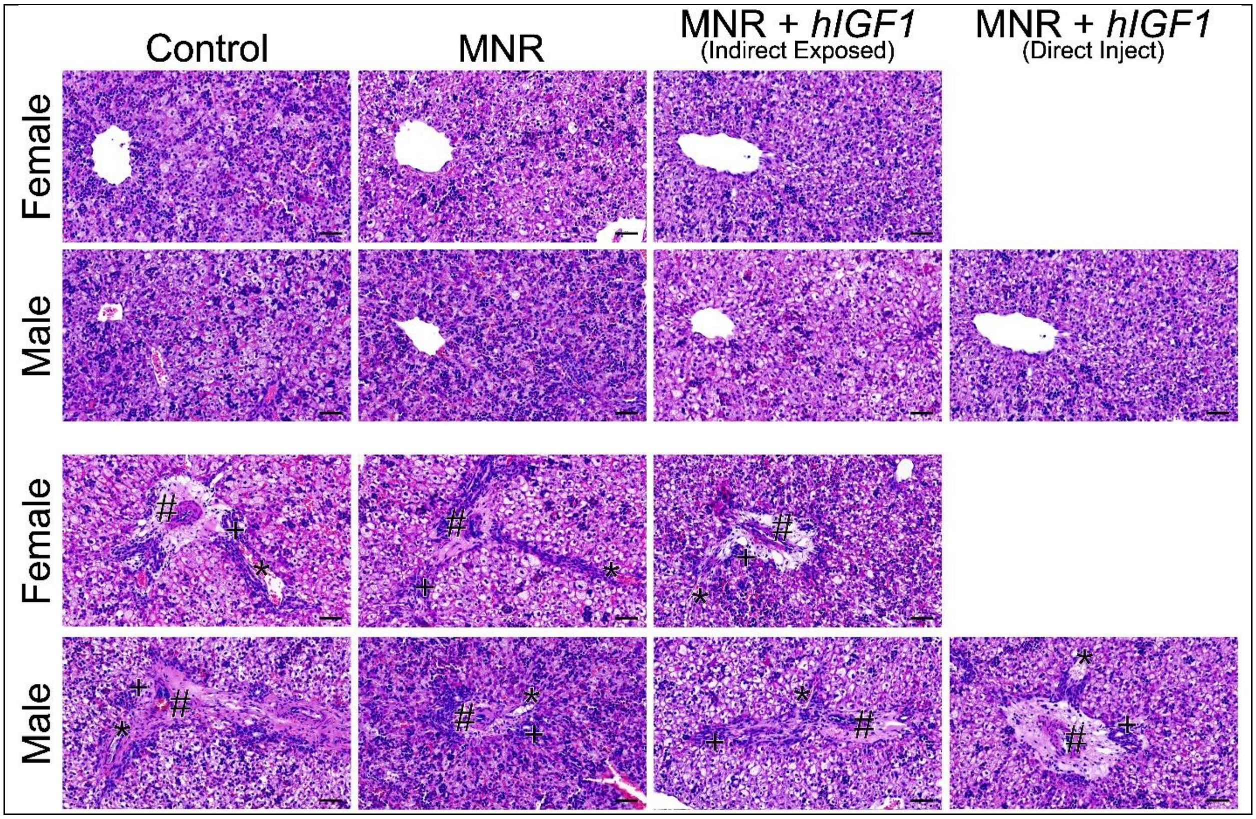

**Supplemental Figure S2. Representative images of hematoxylin and eosin staining in the near-term guinea pig fetal liver.** Assessment of lipid accumulation (upper panels) and liver portal triad morphology (lower panels) and in female and male liver tissue demonstrated very little differences with maternal nutrient restriction (MNR) or *hIGF1* nanoparticle gene therapy treatment. Control and MNR groups were sham treated. Scale bar = 20 µm. # = hepatic artery. * = portal vein. + = bile duct.

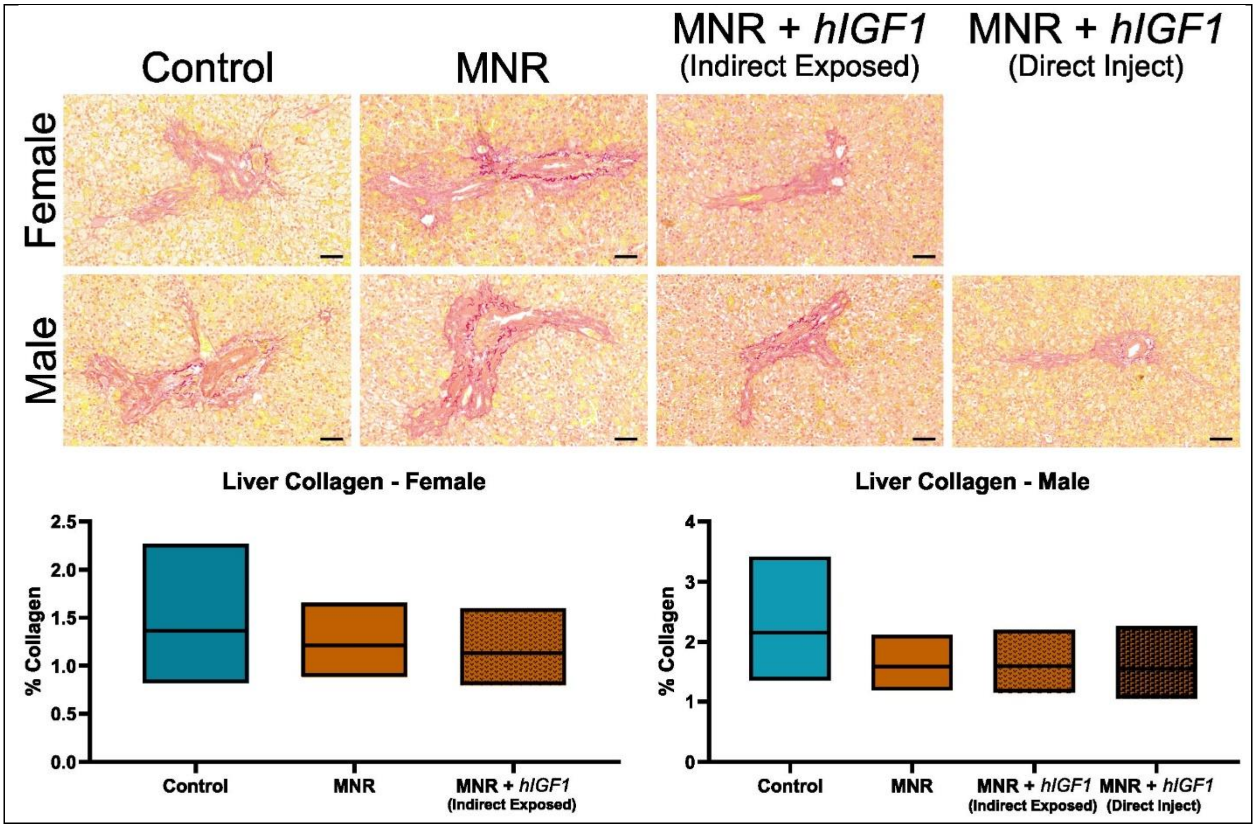


**Supplemental Figure S3. Representative images of collagen deposition (Sirius red stain) around the portal triad of the near-term guinea pig fetal liver.** Quantification of the percentage collagen around the liver portal triads indicated similar levels of collage between sham treated Control, sham treated maternal nutrient restriction (MNR) and MNR *hIGF1* nanoparticle gene therapy treated groups in both female male fetuses. 10 randomly selected portal triads were analyzed using ImageJ Software. Control: n = 6 dams (8 female and 11 male fetuses), MNR: n = 6 dams (5 female and 11 male fetuses), MNR + *hIGF1*: n = 5 dams (6 female and 10 male fetuses). Data are estimated marginal means ± 95% confidence interval. Scale bar = 20 μm

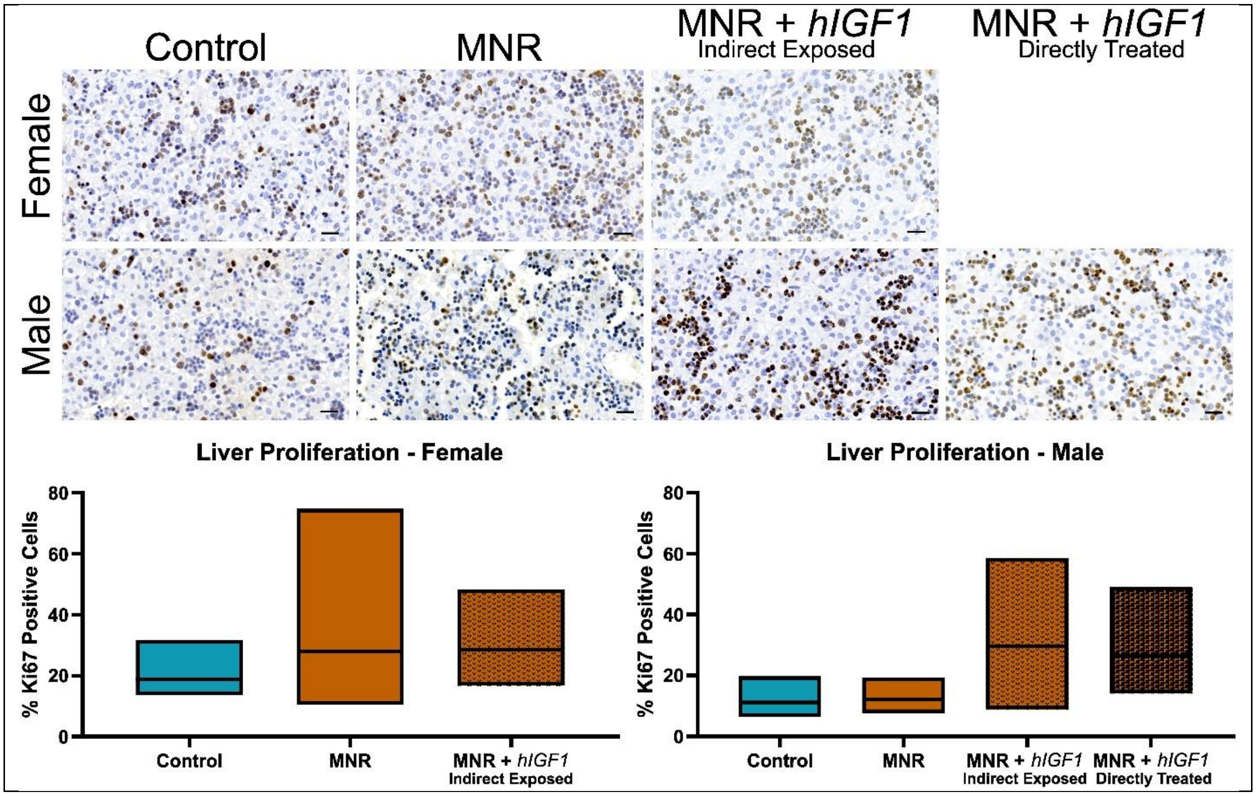


**Supplemental Figure S4. Representative images of hepatocyte proliferation (Ki67 positive nuclei) in the near-term guinea pig fetal liver.** The number of Ki67 positive cells was similar between sham treated Control, sham treated maternal nutrient restriction (MNR) and *hIGF1* nanoparticle gene therapy treated MNR + *hIGF1* groups in female and male fetal livers. 10 randomly selected fields of view were generated and analyzed. Control: n = 6 dams (8 female and 11 male fetuses), MNR: n = 6 dams (5 female and 11 male fetuses), MNR + *hIGF1*: n = 5 dams (6 female and 10 male fetuses). Data are estimated marginal means ± 95% confidence interval. Scale bar = 20 µm.

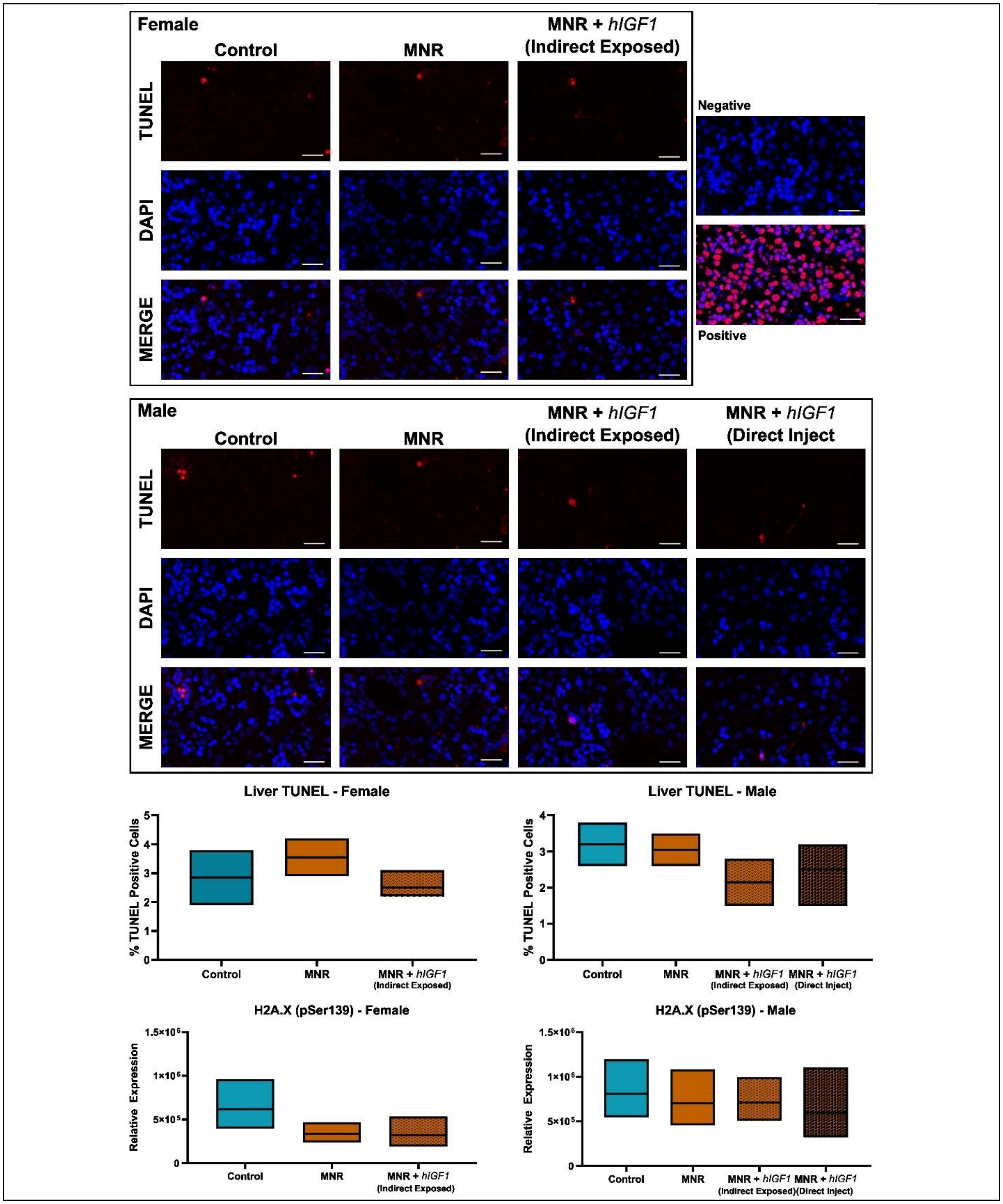


**Supplemental Figure S5. Representative images of apoptosis (TUNEL positive nuclei) and impact on protein expression of H2A.X (pSer139) in the near-term guinea pig fetal liver.** The number of TUNEL positive cells, and protein expression of H2A.X (pSer139), a biomarker of double-stranded DNA damage, was similar between sham treated Control, sham treated maternal nutrient restriction (MNR) and *hIGF1* nanoparticle gene therapy treated MNR + *hIGF1* groups in female and male fetal livers. For TUNEL staining, a negative control excluding the TdT reaction was included. A positive control was generated by incubating tissue with DNase I. 10 randomly selected fields of view were generated and analyzed. For H2A.X (pSer139), representative western blot images are presented in Supplemental Figure S6. Control: n = 6 dams (8 female and 11 male fetuses), MNR: n = 6 dams (5 female and 11 male fetuses), MNR + *hIGF1*: n = 5 dams (6 female and 10 male fetuses). Data are estimated marginal means ± 95% confidence interval. Scale bar = 20 μm.

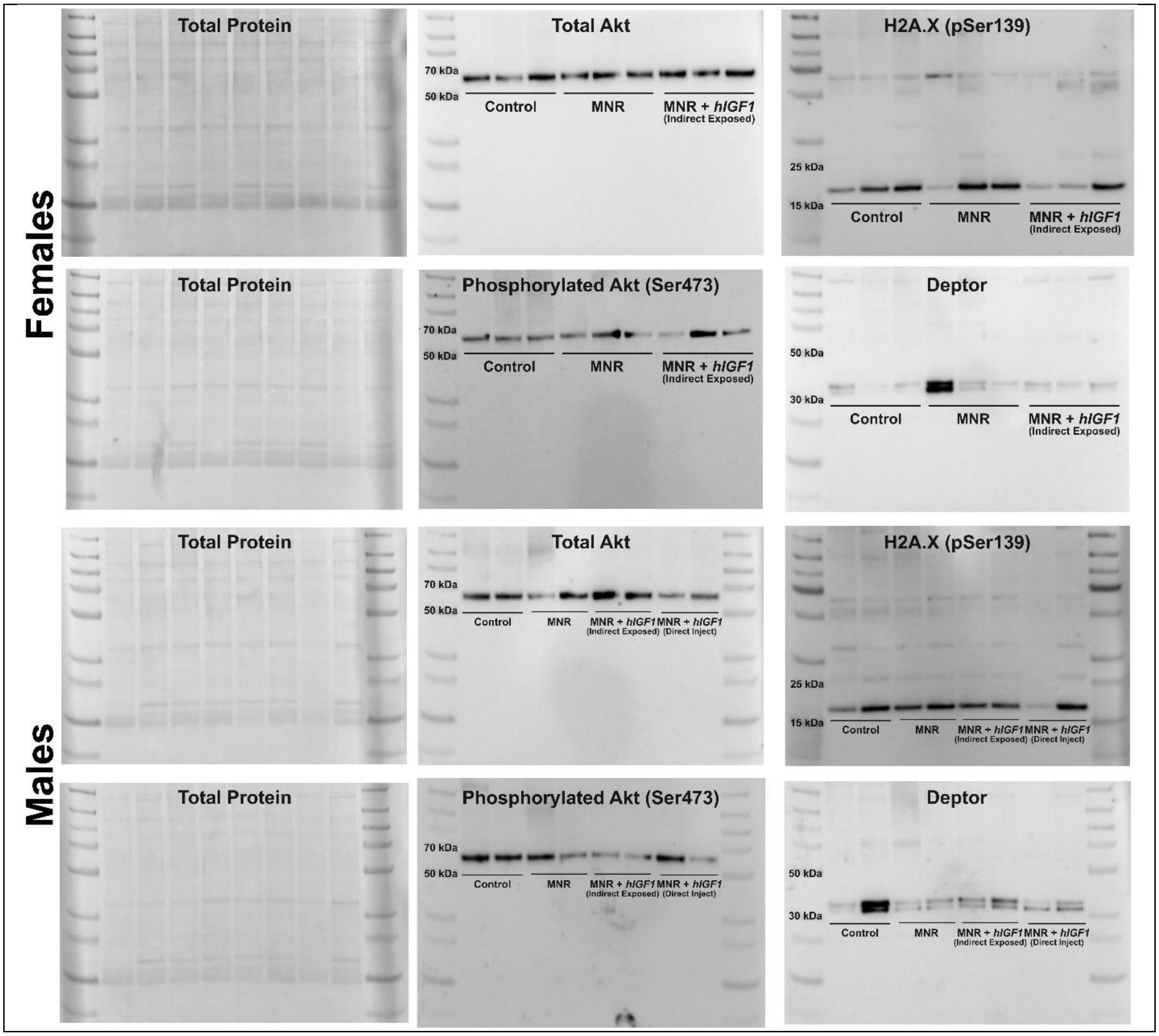


**Supplemental Figure S6. Representative western blot images for analysis of fetal liver protein expression.** Ponceau S staining was used to visualize total protein on nitrocellulose membranes. Membranes containing liver protein from female and male fetuses were first blotted with antibodies to detect Akt and phosphorylated-Akt, imaged, stripped and then re-blotted with antibodies to detect H2A.X (pSer139) and Deptor. Protein abundance was visualized using Chemiluminescence detection on a Biorad Chemidoc, and quantified using ImageLab software

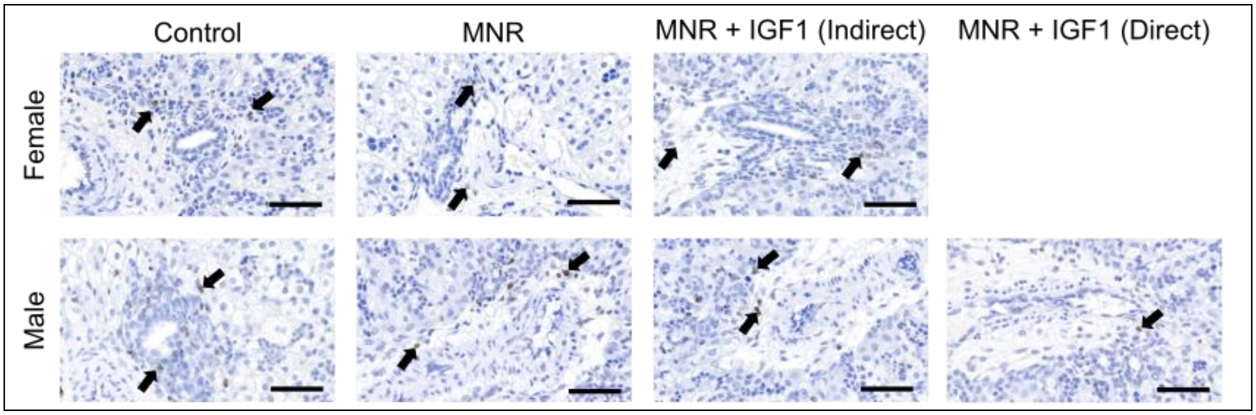


**Supplemental Figure S7. Representative images of CD45 positive immune cells around the portal triad in the near-term guinea pig fetal liver.** Increased immune cell infiltration was not observed around the portal triads with either MNR or placental *hIGF1* nanoparticle treatment in either female or male fetuses. Control: n = 6 dams (8 female and 11 male fetuses), MNR: n = 6 dams (5 female and 11 male fetuses), MNR + *hIGF1*: n = 5 dams (6 female and 10 male fetuses). Scale bar = 20 μm.

**Supplemental Table S1.** Fetal liver expression of lipid and glucose metabolism-related genes

| | Control | | | MNR | | | MNR + *hIGF1* (Indirect Exposed) | | | MNR + *hIGF1* (Directly Treated) | | | P value | | |
|---|---|---|---|---|---|---|---|---|---|---|---|---|---|---|---|
| | | | | | | | | | | | | | Diet | Treatment | Directly Treated (males only) |
| Females | | | | | | | | | | | | | | | |
| | EMM | 95% CI | | EMM | 95% CI | | EMM | 95% CI | | | | | | | |
| Pro-fibrosis Markers | | | | | | | | | | | | | | | |
| *Ctgf* | 1.25 | 0.73 | 2.12 | 1.84 | 1.10 | 3.06 | 2.26 | 1.31 | 3.92 | | | | NS | NS | |
| *Timp1* | 1.16 | 0.92 | 1.46 | 1.17 | 0.91 | 1.51 | 1.76 | 1.11 | 2.80 | | | | NS | NS | |
| *Timp2* | 1.02 | 0.79 | 1.33 | 0.89 | 0.65 | 1.22 | 1.07 | 0.63 | 1.82 | | | | NS | NS | |
| Inflammatory Markers | | | | | | | | | | | | | | | |
| *Tnfa* | 0.93 | 0.53 | 1.61 | 0.70 | 0.36 | 1.36 | 1.22 | 0.31 | 2.85 | | | | NS | NS | |
| *IL6* | 0.74 | 0.53 | 1.03 | 0.61 | 0.37 | 1.00 | 0.96 | 0.77 | 1.20 | | | | NS | NS | |
| *IL6R* | 0.94 | 0.63 | 1.39 | 0.56 | 0.43 | 0.74 | 0.50 | 0.28 | 0.90 | | | | NS | NS | |
| *IL1b* | 1.13 | 0.92 | 1.39 | 0.65 | 0.39 | 1.07 | 0.39 | 0.21 | 0.75 | | | | 0.039 | NS | |
| Glucose Transport and Gluconeogenesis | | | | | | | | | | | | | | | |
| *Pck1* | 1.58 | 1.09 | 2.29 | 0.86 | 0.47 | 1.60 | 1.90 | 0.58 | 6.18 | | | | NS | NS | |
| *GcgR* | 1.39 | 1.12 | 1.72 | 1.44 | 0.81 | 2.56 | 1.47 | 1.06 | 2.04 | | | | NS | NS | |
| *Igf1R* | 1.05 | 0.89 | 1.25 | 1.54 | 0.71 | 3.36 | 0.90 | 0.53 | 1.54 | | | | NS | NS | |
| *Igf2* | 1.53 | 0.93 | 2.49 | 1.09 | 0.78 | 1.53 | 0.89 | 0.44 | 1.81 | | | | NS | NS | |
| *Males* | | | | | | | | | | | | | | | |
| | EMM | 95% CI | | EMM | 95% CI | | EMM | 95% CI | | EMM | 95% CI | | Diet | Treatment | Direct Injection (males only) |
| Pro-fibrosis Markers | | | | | | | | | | | | | | | |
| *Ctgf* | 1.11 | 0.86 | 2.42 | 1.05 | 0.87 | 1.28 | 0.80 | 0.59 | 1.04 | 1.27 | 0.59 | 2.26 | NS | NS | NS |
| *Timp1* | 1.16 | 0.75 | 1.80 | 1.19 | 0.65 | 1.90 | 1.15 | 0.86 | 1.53 | 1.35 | 0.26 | 2.38 | NS | NS | NS |
| *Timp2* | 1.03 | 0.88 | 1.20 | 1.23 | 0.58 | 1.83 | 1.20 | 0.49 | 2.98 | 0.91 | 0.57 | 1.45 | NS | NS | NS |
| Inflammatory Markers | | | | | | | | | | | | | | | |
| *Tnfa* | 1.15 | 0.80 | 1.65 | 1.05 | 0.75 | 1.47 | 1.32 | 0.93 | 1.89 | 1.90 | 1.09 | 3.33 | NS | NS | NS |
| *IL6* | 0.49 | 0.34 | 0.71 | 0.51 | 0.32 | 0.83 | 0.84 | 0.77 | 1.41 | 0.92 | 0.30 | 2.85 | NS | NS | NS |
| *IL6R* | 0.86 | 0.64 | 1.14 | 0.66 | 0.49 | 0.89 | 0.87 | 0.66 | 1.15 | 0.56 | 0.39 | 0.82 | NS | NS | NS |
| *IL1b* | 0.87 | 0.70 | 1.08 | 0.68 | 0.51 | 0.91 | 0.56 | 0.42 | 0.75 | 0.27 | 0.11 | 0.62 | NS | NS | NS |

| | | | | | | | | | | | | | | | |
|---|---|---|---|---|---|---|---|---|---|---|---|---|---|---|---|
| Glucose Transport and Gluconeogenesis | | | | | | | | | | | | | | | |
| *Pck1* | 0.70 | 0.55 | 0.89 | 0.79 | 0.53 | 1.17 | 0.98 | 0.70 | 1.36 | 0.93 | 0.71 | 1.12 | NS | NS | NS |
| *GcgR* | 1.56 | 1.35 | 1.81 | 1.25 | 0.97 | 1.61 | 1.03 | 0.71 | 1.51 | 1.24 | 0.65 | 1.84 | NS | NS | NS |
| *Igf1R* | 1.04 | 0.80 | 1.35 | 1.17 | 0.69 | 1.97 | 1.58 | 1.05 | 2.37 | 1.36 | 0.72 | 2.60 | NS | NS | NS |
| *Igf2* | 2.42 | 1.07 | 5.47 | 1.61 | 1.07 | 2.42 | 1.81 | 1.06 | 3.26 | 1.77 | 0.93 | 3.36 | NS | NS | NS |

CI = Confidence Interval. EMM = Estimated Marginal Mean. MNR = Maternal Nutrient Restriction. Control and MNR are sham treated. Control: n = 6 dams (8 female and 11 male fetuses), MNR: n = 6 dams (5 female and 11 male fetuses), MNR + *hIGF1*: n = 5 dams (6 female and 10 male fetuses). Statistical significance calculated using Generalized Estimating Equations.
